# Maternal and child immune profiles are associated with neurometabolite measures of early-life neuroinflammation in children who are HIV-exposed and uninfected: a South African birth cohort

**DOI:** 10.1101/2025.03.17.643628

**Authors:** Cesc Bertran-Cobo, Frances C Robertson, Tusekile Sarah Kangwa, Jenna Annandale, Sivenesi Subramoney, Katherine L Narr, Shantanu H Joshi, Nadia Hoffman, Heather J Zar, Dan J Stein, Kirsten A Donald, Catherine J Wedderburn, Petrus J W Naudé

**Author notes:** Shared last authorship (Dan J Stein, Kirsten A Donald, Catherine J Wedderburn, Petrus J W Naudé). **Correspondence**: Cesc Bertran-Cobo.

## Abstract

Children who are HIV-exposed and uninfected (HEU) are at risk of neurodevelopmental delays, which may be partially due to maternal immune dysregulation during pregnancy. This study investigates associations between maternal and child immune profiles and early neurometabolite profiles in HEU and HIV-unexposed (HU) children from a South African birth cohort. A subgroup of 156 children (66 HEU, 90 HU) from the Drakenstein Child Health Study underwent magnetic resonance spectroscopy at age 2–3 years, and maternal and child serum markers were measured at multiple timepoints via immunoassays.

In HEU children, serum concentrations of maternal pro-inflammatory cytokines IL-5 (β=0.79, p=0.005) and IL-8 (β=0.64, p=0.02) were associated with myo-inositol ratios in parietal grey and white matter regions, respectively, while child serum MMP-9 at two years was associated with myo-inositol ratios in the midline parietal grey matter (β=1.30, p=0.03). The association of maternal anti-inflammatory cytokine IL-13 with glutamate ratios in the midline parietal grey matter was negative in HEU (β=-0.41, p=0.038) and positive in HU children (β=0.42, p<0.0001). These findings suggest maternal immune activation may affect neurometabolite profiles in HEU children.

## Introduction

An estimated 39 million people live with HIV worldwide, with approximately 20.8 million residing in Southern and Eastern Africa (1). Widespread access to antiretroviral therapy (ART) and vertical transmission prevention programmes have significantly reduced new HIV infections (1). As a result, the number of new infections among children under 14 years has decreased by over 60% since 2000 (2). This has led to the emergence of a rapidly growing population of children who are born to mothers living with HIV and remain uninfected themselves (HIV-exposed and uninfected [HEU]) (3,4). Children who are HEU are estimated at 16.1 million globally (1), with the highest number residing in South Africa (2). This group, while spared from HIV infection, faces unique health challenges, including increased mortality rates (5) and a higher risk of adverse neurodevelopmental outcomes compared to children who are HIV-unexposed (HU) (6).

Prenatal exposure to maternal HIV and ART may alter fetal immune programming, resulting in distinct immune profiles in children who are HEU compared to their HU peers (7,8). However, findings are mixed within and across geographical regions. Studies in Southern African cohorts have shown decreased pro- and anti-inflammatory cytokine levels in HEU children (9), while others found no differences (10). In contrast, Latin American studies reported elevated levels of inflammatory and monocyte activation markers in HEU compared to HU children (11,12). These inconsistencies highlight the need for further research to elucidate how HIV exposure affects immune and neurodevelopmental trajectories in children across different social and geographical contexts (7).

In HEU infants from a South African birth cohort, the Drakenstein Child Health Study (DCHS), increased serum inflammatory cytokines at 6 weeks of age predicted poorer motor and language development at two years (10). One proposed pathway linking maternal HIV to child immune profiles and neurodevelopmental outcomes is through the effect of maternal inflammation on the developing fetal brain. Pregnant women living with HIV may experience chronic immune activation despite ART (13,14), resulting in altered cytokine levels that can cross the placenta and impact glial cell integrity (15,16) and white matter development (17,18). Maternal immune activation has been associated with altered brain growth and connectivity in the offspring, potentially increasing the risk for neurodevelopmental disorders (19–21). It has been hypothesized that prenatal exposure to maternal immune activation can predispose children who are HEU to exaggerated neuroinflammatory responses when exposed to subsequent postnatal insults, potentially disrupting typical brain development (18,22).

Given the critical role of the immune system in shaping brain development, a deeper understanding of how immune profiles in children who are HEU change over time and their potential impact on brain health is needed. Magnetic resonance spectroscopy (MRS) provides a clinically established method for *in vivo* quantification of key brain metabolites such as myo-inositol and glutamate, which are linked to neuroinflammation and neuronal function, respectively (23,24). Our research found elevated myo-inositol ratios to creatine in the parietal white matter of South African HEU children at 2–3 years of age, suggesting a neuroinflammatory response due to maternal HIV status (25). Other studies in older children have also reported neurometabolite differences between HEU and HU groups (26–28). Taken together, these findings suggest that HIV-associated changes to the maternal immune system may contribute to altered child neurometabolite patterns, potentially leading to neurodevelopmental delays observed in HEU children. To date, no studies have examined longitudinal changes in serum marker levels through childhood nor their potential associations with neuroimaging findings in children who are HEU. This approach is essential to delineate the temporal dynamics of immune profiles in this population and their relationship to neurobiological outcomes.

Our study aimed to address this gap by investigating maternal and child peripheral blood immune profiles from pregnancy through birth and early childhood, and their associations with brain metabolite levels at age 2–3 years, in a cohort of South African HEU and HU children from the DCHS (29). We hypothesized that mothers living with HIV and their HEU children would show altered levels of serum markers compared to mothers without HIV and their HU children, and that these alterations would be associated with child elevated levels of myo-inositol in the parietal white matter at age 2–3 years.

## Results

### Cohort and demographic characteristics

Out of the 1143 mother-child pairs enrolled in the DCHS, a group of 156 children underwent MRS imaging at age 2–3 years. Data was successfully acquired from 143 children in the parietal grey matter, 134 in the left parietal white matter, and 92 in the right parietal white matter. Following quality assessment, 9 participants were excluded due to low-quality spectra, resulting in a subset of 83 children (36 HEU, 47 HU) with high-quality data available for all three parietal brain regions.

Sociodemographic characteristics were generally comparable between mothers living with HIV (n=66) and those without HIV (n=90) in this cohort, with no significant differences in household income, education, employment, marital status, or most perinatal health behaviors (p>0.05). However, mothers living with HIV were older at delivery, had lower rates of depression and tobacco use during pregnancy, were less likely to be single mothers, and breastfed for a shorter duration than mothers not living with HIV. Children in HEU and HU groups had comparable birth weights and nutritional statuses (**Table 1**).

**Table 1.**
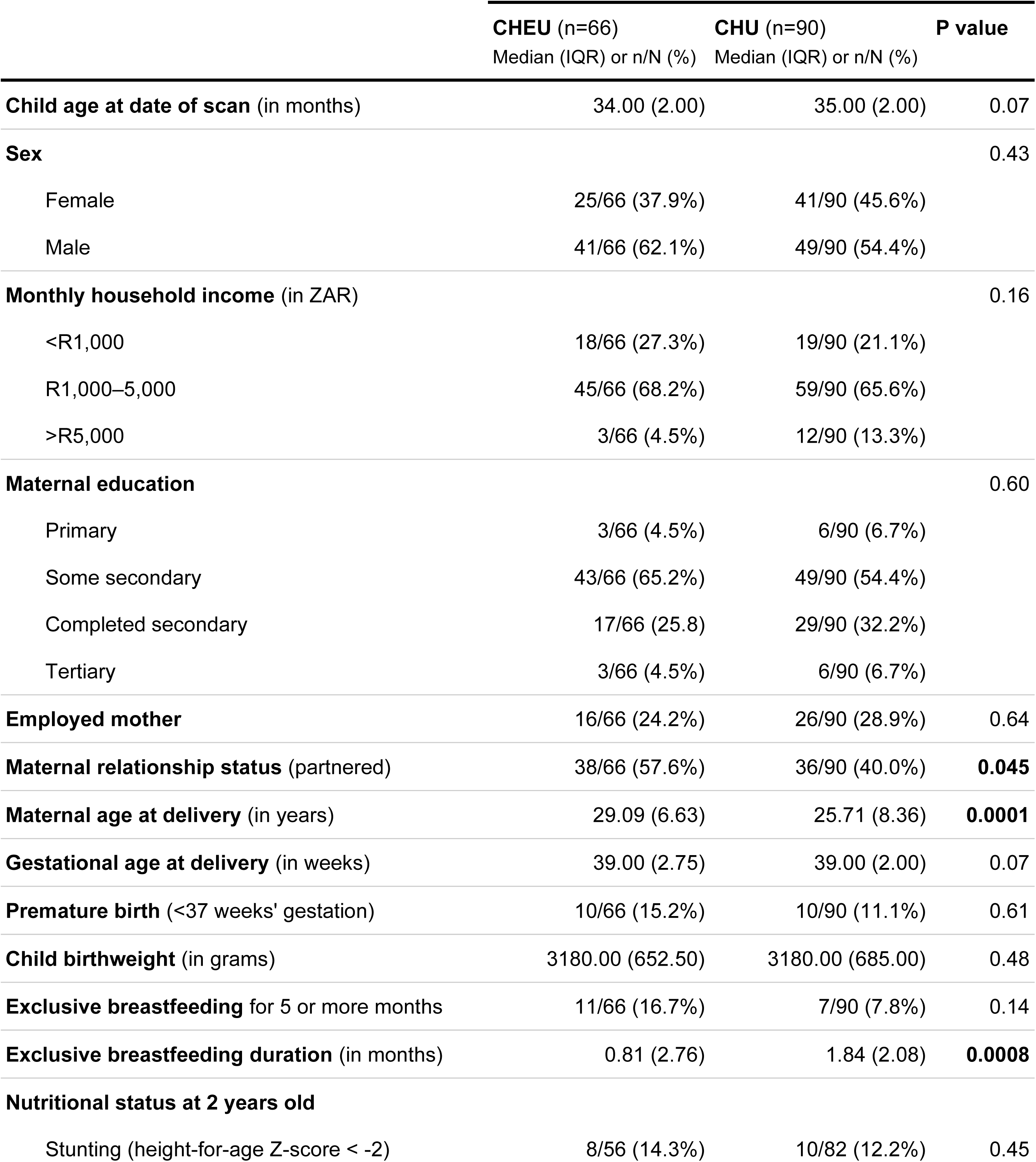
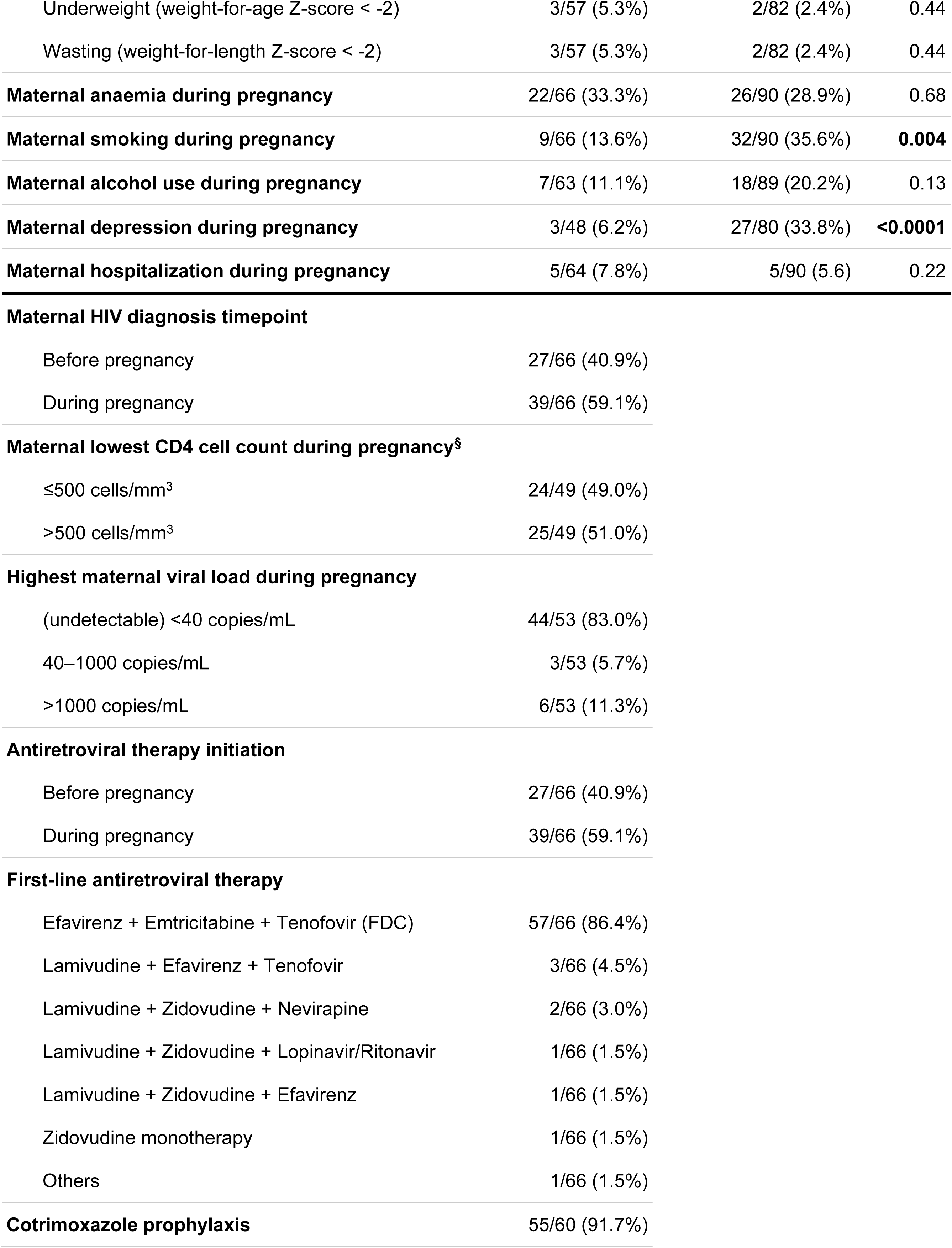
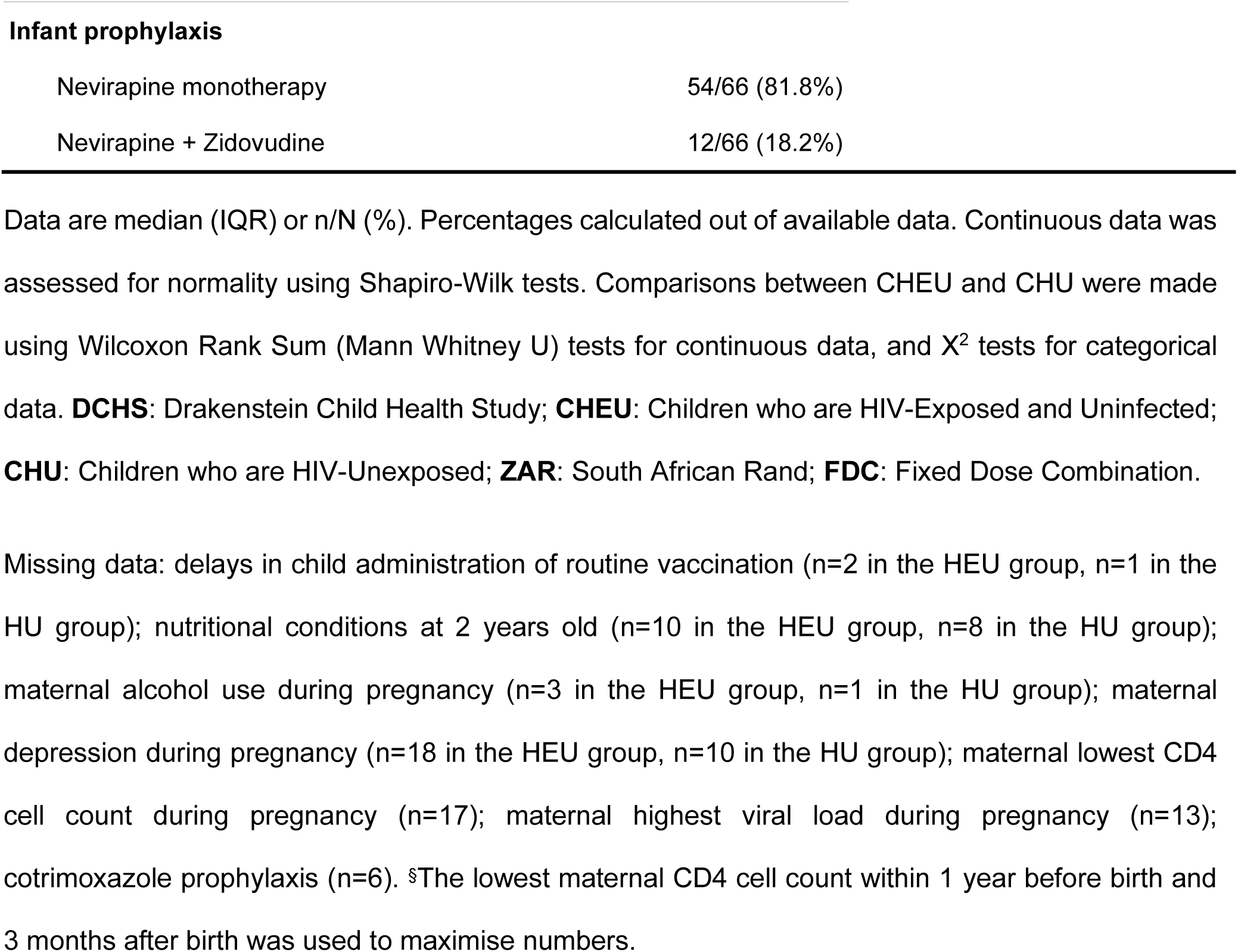
Sociodemographic characteristics of children invited for MRS at age 2–3 years, according to HIV exposure.

|  | **CHEU (n=66)**<br>Median (IQR) or n/N (%) | **CHU (n=90)**<br>Median (IQR) or n/N (%) | **P value** |
| --- | --- | --- | --- |
| **Child age at date of scan** (in months) | 34.00 (2.00) | 35.00 (2.00) | 0.07 |
| **Sex** |  |  | 0.43 |
| Female | 25/66 (37.9%) | 41/90 (45.6%) |  |
| Male | 41/66 (62.1%) | 49/90 (54.4%) |  |
| **Monthly household income** (in ZAR) |  |  | 0.16 |
| <R1,000 | 18/66 (27.3%) | 19/90 (21.1%) |  |
| R1,000–5,000 | 45/66 (68.2%) | 59/90 (65.6%) |  |
| >R5,000 | 3/66 (4.5%) | 12/90 (13.3%) |  |
| **Maternal education** |  |  | 0.60 |
| Primary | 3/66 (4.5%) | 6/90 (6.7%) |  |
| Some secondary | 43/66 (65.2%) | 49/90 (54.4%) |  |
| Completed secondary | 17/66 (25.8) | 29/90 (32.2%) |  |
| Tertiary | 3/66 (4.5%) | 6/90 (6.7%) |  |
| **Employed mother** | 16/66 (24.2%) | 26/90 (28.9%) | 0.64 |
| **Maternal relationship status** (partnered) | 38/66 (57.6%) | 36/90 (40.0%) | **0.045** |
| **Maternal age at delivery** (in years) | 29.09 (6.63) | 25.71 (8.36) | **0.0001** |
| **Gestational age at delivery** (in weeks) | 39.00 (2.75) | 39.00 (2.00) | 0.07 |
| **Premature birth** (<37 weeks' gestation) | 10/66 (15.2%) | 10/90 (11.1%) | 0.61 |
| **Child birthweight** (in grams) | 3180.00 (652.50) | 3180.00 (685.00) | 0.48 |
| **Exclusive breastfeeding** for 5 or more months | 11/66 (16.7%) | 7/90 (7.8%) | 0.14 |
| **Exclusive breastfeeding duration** (in months) | 0.81 (2.76) | 1.84 (2.08) | **0.0008** |
| **Nutritional status at 2 years old** |  |  |  |
| Stunting (height-for-age Z-score < -2) | 8/56 (14.3%) | 10/82 (12.2%) | 0.45 |
| Underweight (weight-for-age Z-score < -2) | 3/57 (5.3%) | 2/82 (2.4%) | 0.44 |
| Wasting (weight-for-length Z-score < -2) | 3/57 (5.3%) | 2/82 (2.4%) | 0.44 |
| **Maternal anaemia during pregnancy** | 22/66 (33.3%) | 26/90 (28.9%) | 0.68 |
| **Maternal smoking during pregnancy** | 9/66 (13.6%) | 32/90 (35.6%) | **0.004** |
| **Maternal alcohol use during pregnancy** | 7/63 (11.1%) | 18/89 (20.2%) | 0.13 |
| **Maternal depression during pregnancy** | 3/48 (6.2%) | 27/80 (33.8%) | **<0.0001** |
| **Maternal hospitalization during pregnancy** | 5/64 (7.8%) | 5/90 (5.6) | 0.22 |
| **Maternal HIV diagnosis timepoint** |  |  |  |
| Before pregnancy | 27/66 (40.9%) |  |  |
| During pregnancy | 39/66 (59.1%) |  |  |
| **Maternal lowest CD4 cell count during pregnancy<sup>s</sup>** |  |  |  |
| ≤500 cells/mm<sup>3</sup> | 24/49 (49.0%) |  |  |
| >500 cells/mm<sup>3</sup> | 25/49 (51.0%) |  |  |
| **Highest maternal viral load during pregnancy** |  |  |  |
| (undetectable) <40 copies/mL | 44/53 (83.0%) |  |  |
| 40–1000 copies/mL | 3/53 (5.7%) |  |  |
| >1000 copies/mL | 6/53 (11.3%) |  |  |
| **Antiretroviral therapy initiation** |  |  |  |
| Before pregnancy | 27/66 (40.9%) |  |  |
| During pregnancy | 39/66 (59.1%) |  |  |
| **First-line antiretroviral therapy** |  |  |  |
| Efavirenz + Emtricitabine + Tenofovir (FDC) | 57/66 (86.4%) |  |  |
| Lamivudine + Efavirenz + Tenofovir | 3/66 (4.5%) |  |  |
| Lamivudine + Zidovudine + Nevirapine | 2/66 (3.0%) |  |  |
| Lamivudine + Zidovudine + Lopinavir/Ritonavir | 1/66 (1.5%) |  |  |
| Lamivudine + Zidovudine + Efavirenz | 1/66 (1.5%) |  |  |
| Zidovudine monotherapy | 1/66 (1.5%) |  |  |
| Others | 1/66 (1.5%) |  |  |
| **Cotrimoxazole prophylaxis** | 55/60 (91.7%) |  |  |

**Infant prophylaxis**
|  |  |
| --- | --- |
| Nevirapine monotherapy | 54/66 (81.8%) |
| Nevirapine + Zidovudine | 12/66 (18.2%) |

Data are median (IQR) or n/N (%). Percentages calculated out of available data. Continuous data was assessed for normality using Shapiro-Wilk tests. Comparisons between CHEU and CHU were made using Wilcoxon Rank Sum (Mann Whitney U) tests for continuous data, and $X^2$ tests for categorical data. **DCHS**: Drakenstein Child Health Study; **CHEU**: Children who are HIV-Exposed and Uninfected; **CHU**: Children who are HIV-Unexposed; **ZAR**: South African Rand; **FDC**: Fixed Dose Combination.
Missing data: delays in child administration of routine vaccination (n=2 in the HEU group, n=1 in the HU group); nutritional conditions at 2 years old (n=10 in the HEU group, n=8 in the HU group); maternal alcohol use during pregnancy (n=3 in the HEU group, n=1 in the HU group); maternal depression during pregnancy (n=18 in the HEU group, n=10 in the HU group); maternal lowest CD4 cell count during pregnancy (n=17); maternal highest viral load during pregnancy (n=13); cotrimoxazole prophylaxis (n=6). <sup>§</sup>The lowest maternal CD4 cell count within 1 year before birth and 3 months after birth was used to maximise numbers.

All mothers living with HIV received ART, either before conception (40.9%) or during pregnancy (59.1%). Most (86.4%) were on a first-line regimen of efavirenz, emtricitabine, and tenofovir, and nearly all (91.7%) received cotrimoxazole prophylaxis. All HEU children received post-exposure prophylaxis, either nevirapine alone (81.8%) or in combination with zidovudine (18.2%) (**Table 1**). In the subset of 83 children with complete MRS data, the only significant differences between HEU and HU groups were in maternal age at delivery and antenatal depression (**Supplementary Table 2A**). Overall, the complete MRS subset of participants was comparable to the initial group of 156 children (**Supplementary Table 2B**), ensuring representativeness for subsequent analyses.

### Group differences in maternal and child serum marker concentrations

#### Maternal markers during pregnancy

Out of 156 mothers whose children underwent MRS at age 2–3 years, 138 (88.5%, n=60 with HIV, n=78 not with HIV) had available serum marker data during pregnancy (**Supplementary Figure 1A**). Several differences were observed based on HIV status. Levels of pro-inflammatory cytokine GM-CSF and neuroinflammatory marker MMP-9 were significantly lower in mothers living with HIV compared to their peers, remaining after Benjamini-Hochberg (BH) adjustment for multiple comparisons (GM-CSF corrected p=0.034; MMP-9 corrected p=0.002) (**Supplementary Figure 2**, **Supplementary Table 3**). Prior to BH correction, mothers living with HIV were found to have lower circulating levels of the neuroinflammatory marker NGAL (uncorrected p=0.018) and higher levels of the monocyte activation marker CD14 (uncorrected p= 0.044) in comparison to their counterparts. For the remaining markers, no significant group differences were found.

#### Infant markers at 6 weeks of age

97/156 infants (62.2%, n=41 HEU, n=56 HU) had available serum marker measurements at 6 weeks post-birth (**Supplementary Figure 1A**). NGAL levels were significantly lower in HEU compared to HU infants in the unadjusted analysis (p= 0.032), but this finding did not survive BH correction for multiple comparisons (**Supplementary Figure 2**, **Supplementary Table 3**).

#### Child markers at 2 years of age

In the 2-year timepoint, 111/156 children (71.2%, n=46 HEU, n=65 HU) had available serum marker data (**Supplementary Figure 1A**). Analyses indicated that levels of pro-inflammatory cytokine IL-1β (uncorrected p=0.032) and anti-inflammatory cytokine IL-12p70 (uncorrected p=0.048) were lower in HEU children compared to HU, but these differences did not remain significant after BH adjustment (**Supplementary Figure 2**, **Supplementary Table 3**).

### Child serum marker trajectories

We used Linear Mixed-effects Models (LMMs) to examine longitudinal changes in circulating marker levels between ages 6 weeks and 2 years in 127/156 children (81.4%, n=56 HEU, n=71 HU) with available serum marker data at either of the two timepoints. To address the issue of missing data, Maximum Likelihood Estimation was applied in our LMMs, with use of the *lme4* package in R. Akaike Information Criterion (AIC) and Bayesian Information Criterion (BIC) values revealed that, out of three covariance structures tested, the model with variance components fitted our data better than the compound symmetry and unstructured covariance models.

No significant longitudinal differences were found between 6 weeks and 2 years of age in our group of 127 participants or between HEU and HU subgroups (**Supplementary Figure 3**, **Supplementary Table 4**).

### Cross-sectional associations between serum markers and brain metabolite ratios

Linear models were run in a subset of 83 children (36 HEU, 47 HU) with high-quality MRS data for all three parietal brain regions. Adjusted analyses revealed a significant influence of tissue composition on total choline ratios across all parietal voxels, therefore, this metabolite was excluded from all analyses. N-acetyl-aspartate ratios in the left parietal white matter were also excluded from our analyses due to suboptimal tissue composition.

Results that were significant in unadjusted linear models, survived BH correction, and remained significant after adjusting for potential confounders are reported in **Table 2**. For a full overview of all results, see **Supplementary Figure 4** and **Supplementary Table 5**, as well as **Supplementary Table 6** for subsequent sensitivity analyses.

**Table 2.**
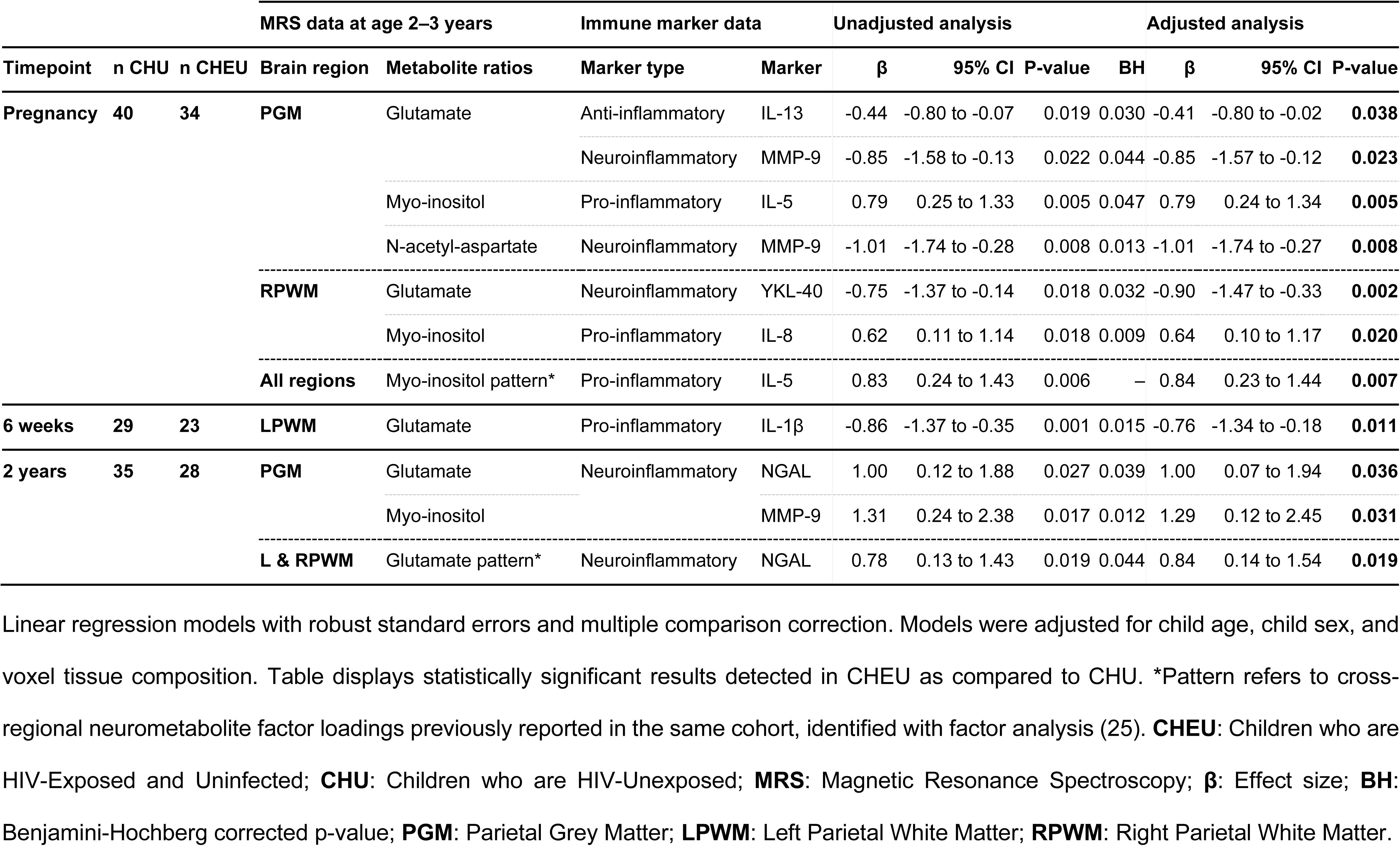
Associations between maternal and child immune marker concentrations and child neurometabolite ratios.

### Maternal serum marker levels and child neurometabolite ratios

Of the 83 children with high-quality MRS data for all three parietal brain regions, 74 (89.2%, n=34 HEU, n=40 HU) had available data on maternal blood marker concentrations during pregnancy (**Supplementary Figure 1B**).

#### Myo-inositol

Maternal IL-5 levels during pregnancy were found to be significantly associated with myo-inositol ratios in the parietal grey matter of children who are HEU (β=0.79 p=0.0053), and maternal IL-8 was found to be associated with myo-inositol ratios in the right parietal white matter of the same group (β=0.64, p=0.0203). Maternal IL-5 was also found to be associated with the composite factor reflecting a pattern of elevated myo-inositol across voxels in children who are HEU (β=0.84, p=0.0075) (**Figure 1**).

**Figure 1.**
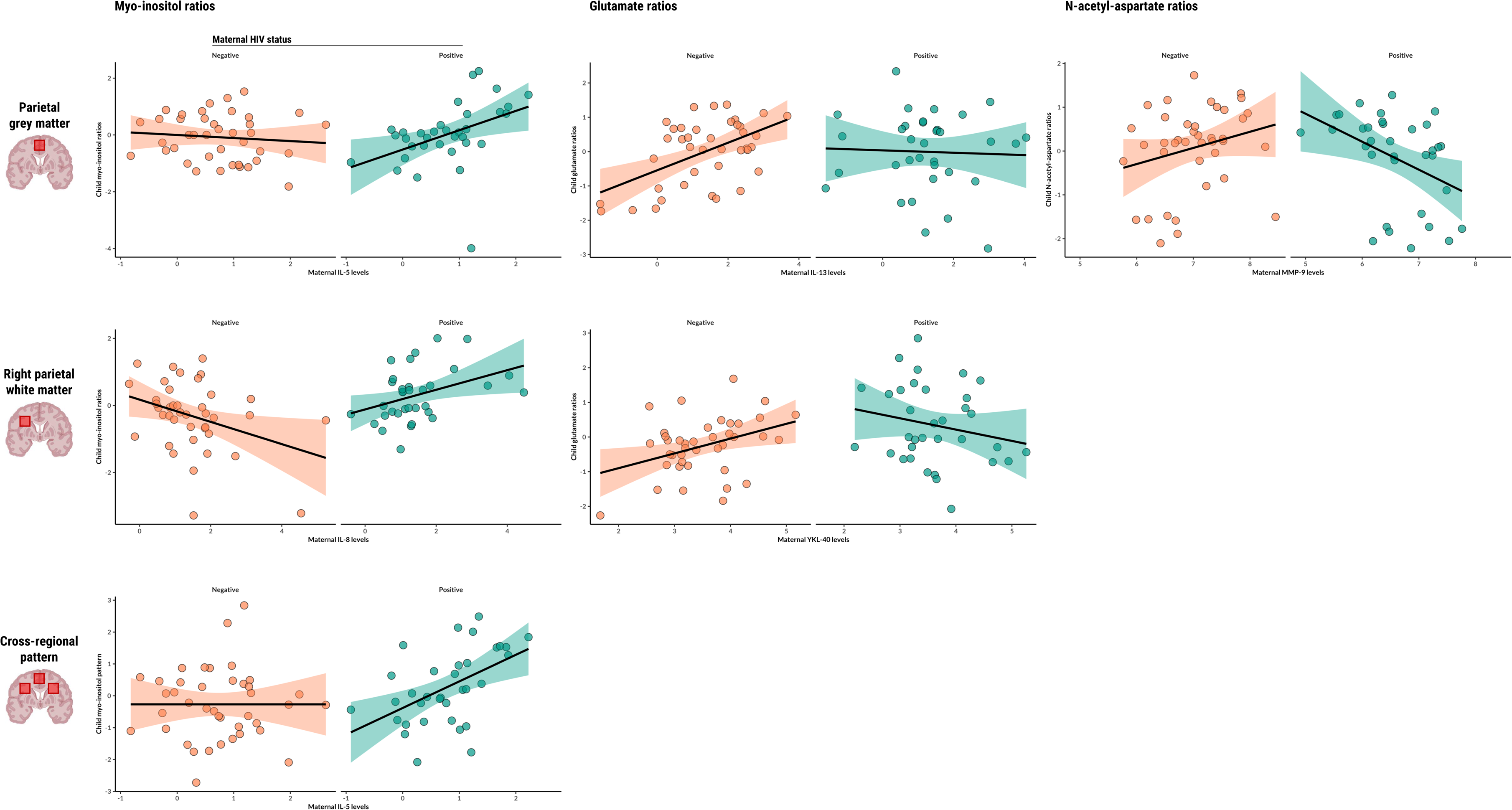
Associations between maternal or child serum markers and child neurometabolite ratios. Scatter plots displaying statistically significant associations between maternal or child serum markers (IL-5, IL-8, IL-13, MMP-9, YKL-40) and child neurometabolite ratios to total creatine (myo-inositol, glutamate, N-acetyl-aspartate) in specific parietal brain regions, as found in adjusted linear regression analyses. Regression lines represent the direction and strength of the associations, with 95% confidence intervals shaded. Data points are colour-coded based on maternal HIV status.

#### Glutamate

Maternal IL-13 levels during pregnancy were positively associated with glutamate ratios in the parietal grey matter of HU children (β=0.42, p<0.0001), and this relationship was significantly modified by maternal HIV infection (β=-0.41, p=0.0380). In the same brain region, maternal MMP-9 levels showed an association with glutamate ratios of children who are HEU (β=-0.85, p=0.0231). Additionally, maternal YKL-40 was positively associated with child glutamate ratios in the right parietal white matter of HU children (β=0.52, p=0.0046) and maternal HIV infection modified this relationship (β=-0.90, p=0.0024). Child sex was identified as a significant covariate in right parietal white matter analyses.

#### N-acetyl-aspartate

Maternal MMP-9 levels during pregnancy were found to be associated with N-acetyl-aspartate ratios in the parietal grey matter of children who are HEU (β=-1.01, p=0.0081).

##### Infant serum marker levels and child neurometabolite ratios

Data on blood marker concentrations at 6 weeks of age were available for 52 out of 83 children (62.7%, n=23 HEU, n=29 HU) (**Supplementary Figure 1B**). Infant IL-1β levels were found to be associated with glutamate ratios in the left parietal white matter of HU children at age 2–3 years (β=0.28, p=0.0259), and maternal HIV infection modified this relationship (β=-0.76, p=0.0113).

##### Child serum marker levels and neurometabolite ratios at age 2–3 years

Serum marker data at 2 years of age were available for 63 out of 83 participants (75.9%, n=28 HEU, n=35 HU) (**Supplementary Figure 1B**). In the parietal grey matter, levels of the neuroinflammatory marker MMP-9 showed an association with myo-inositol ratios of children who are HEU (β=1.29, p=0.0308). The same was found for NGAL levels and glutamate ratios in the HIV-exposed group (β=1.00, p=0.0364). Additionally, NGAL was also found to be associated with the composite factor reflecting a bilateral white matter pattern of glutamate in the HEU group (β=0.84, p=0.0193).

### Mediation effects

Mediation analyses using structural equation modelling were conducted to investigate whether serum markers found to display significant associations with neurometabolites (**Table 2**) would mediate the relationship between maternal HIV infection and child brain metabolite ratios at 2 years of age. A series of models were evaluated, each assessing a distinct serum marker-neurometabolite pair (**Supplementary Table 7**).

Across all models, goodness-of-fit indices consistently indicated a good fit to the data. However, neither the direct paths between maternal HIV infection and child neurometabolite ratios nor the indirect effects through candidate markers were statistically significant. This was consistent across all models (**Supplementary Table 7**).

## Discussion

In this study, we investigated the associations of maternal HIV status with maternal and child serum markers and brain metabolite ratios to creatine in a South African cohort of young children. We found, firstly, differences in immune marker concentrations during pregnancy by maternal HIV status and, secondly, associations between maternal, infant, and child serum markers with child neurometabolite ratios in parietal brain regions at age 2–3 years that differed by HIV exposure. These findings suggest potential pathways through which maternal HIV status may influence early brain development in children and provide insights into how *in utero* changes to maternal immune markers may shape early neurometabolite profiles, potentially through alterations in maternal and child immune systems.

Mothers living with HIV in our cohort had lower levels of the neuroinflammatory marker MMP-9 and pro-inflammatory cytokine GM-CSF compared to mothers not living with HIV, consistent with findings from a larger sample of the same cohort (9). Similarly, decreased MMP-9 levels have been reported in adults living with HIV in other South African settings (30), suggesting a broader trend across populations. While differences in infant and child serum marker levels did not persist after correction for multiple comparisons, we observed lower NGAL levels in HEU infants and lower IL-1β and IL-12p70 in HEU children at 2 years of age before correction. This may indicate that some serum marker differences present at birth diminish over time, whereas others emerge later in childhood. Consistent with this, a recent study on the effects of maternal inflammation on newborn serum profiles found that by six months of age, many HEU serum markers that initially differed had normalized to levels comparable to HU infants (31). Despite this, our longitudinal analysis from six weeks to two years did not reveal significant differences in serum marker trajectories between HEU and HU children.

Our association analyses revealed interactions between maternal HIV status, serum marker concentrations, and child myo-inositol ratios. In children who are HEU, increases in maternal pro-inflammatory cytokines IL-5 and IL-8 were associated with higher child myo-inositol ratios in the parietal grey and white matter regions, respectively. A similar pattern was observed for child MMP-9 levels at age 2 years and myo-inositol ratios in the parietal grey matter at the same timepoint, supporting our hypothesis that maternal and child immune markers are associated with early-life elevated myo-inositol in the parietal white matter of children who are HEU. These results build on our previous work, which identified group differences in myo-inositol ratios in the parietal white matter of this cohort, as well as an association between maternal HIV status and a cross-regional pattern of elevated myo-inositol (25). Clinically, myo-inositol is recognized as a marker of neuroinflammation and gliosis (23,24); thus, increases in myo-inositol may potentially result from maternal immune dysregulation during pregnancy. Inflammatory patterns of myo-inositol have also been linked to cognitive impairment in children living with HIV (26,32).

Maternal levels of anti-inflammatory IL-13 were positively associated with child glutamate ratios in the parietal grey matter of HU children; however, the association was not observed in the HEU group. In contrast, levels of neuroinflammatory MMP-9 in mothers living with HIV were negatively associated with child ratios of this neurotransmitter in the same brain region. Further, infant pro-inflammatory IL-1β at 6 weeks, and neuroinflammatory NGAL at 2 years, were both associated with glutamate ratios in the parietal grey matter of HEU children. Glutamate is a key excitatory neurotransmitter, essential for synaptic plasticity and neurodevelopment (23,24). Its regulation is influenced by astrocytes, which help maintain glutamate balance and prevent excitotoxicity (33). It has been reported that the anti-inflammatory cytokine IL-13 may support astrocyte function and promote cognitive processes by fostering synaptic plasticity and neuroprotection (34,35). HEU children in our study may therefore be at risk of disrupted IL-13-associated regulation of glutamate, potentially contributing to impaired synaptic plasticity and increased vulnerability to neuroinflammatory processes. This disruption could be a potential pathway to explain the neurodevelopmental delays observed in HEU children (6). The relationship between marker concentrations and child neurometabolite ratios was not uniform across HEU and HU groups. It is plausible that maternal immune dysregulation due to HIV creates a unique prenatal environment that sensitizes the fetal brain to subsequent neuroimmune challenges.

Despite the robust associations observed between serum markers and neurometabolites, our mediation analyses did not identify any markers that significantly mediated the relationship between maternal HIV and child brain metabolite profiles at 2–3 years. This suggests that while certain markers are altered in mothers living with HIV, and are associated with brain metabolite patterns, they do not fully account for the impact of maternal HIV exposure on child brain development. The mediatory role of maternal immune activation on child brain health across diverse infection and exposure contexts has been recently reviewed (36). In the context of typical child development, maternal immune activation —measured via circulating IL-6 during pregnancy— was found to mediate neurometabolite ratios and motor development in newborns (37). In our study, maternal IL-6 was associated with child myo-inositol ratios in the right parietal white matter voxel prior to multiple comparison correction. These differences highlight the variability in how maternal immune dysregulation may influence child brain development and suggest that pathways beyond immune markers, such as antiretroviral drug exposure (38,39) or other environmental and psychosocial factors (40,41), may also contribute to the observed neurometabolite alterations in HEU children.

The main strengths of this study include the use of a well-characterized South African cohort and the comprehensive assessment of maternal and child immune profiles at multiple timepoints, combined with high-quality MRS data. Our findings provide a nuanced understanding of how maternal HIV infection affects maternal and child immune profiles and their relationship to early brain development. However, several limitations should be noted. First, although our sample size was larger than those in many previous studies, it was relatively small for some subgroup analyses, which may have limited the power to detect smaller effects. And second, residual confounding cannot be ruled out, particularly given the complex interplay between immune, neurobiological, and environmental factors. We conducted sensitivity analyses to account for potential confounders, including maternal depression and alcohol use during pregnancy as both have been associated with poorer cognitive outcomes in children who are HEU (40,41), and found that our results held.

Future studies will focus on delineating the temporal evolution of neurometabolite and immune profiles in children who are HEU, integrating neurodevelopmental assessments to capture the potential clinical impacts of prenatal HIV exposure. Investigating the combined effects of maternal immune activation, ART regimens, and psychosocial factors may help identify the underlying mechanisms driving the observed neurometabolite alterations. Such insights could inform targeted interventions to mitigate neurodevelopmental risks in HEU children.

In conclusion, this study demonstrates that maternal HIV status is associated with distinct alterations in maternal and child immune profiles, which in turn influence neurometabolite patterns in early childhood. Our findings suggest that perinatal HIV exposure and its immunological impact may create a unique risk profile for HEU children, emphasizing the need for optimal HIV management in pregnancy to mitigate immune dysregulation. Further research is needed to elucidate neuroinflammatory patterns in HEU children and their long-term consequences for neurodevelopment. Finally, these findings highlight the need for strategies that support optimal neurodevelopment in this vulnerable population.

## Methods

### Participants

This study included a sub-sample of mothers and their children from the DCHS, a population-based birth cohort study conducted in a peri-urban area of the Western Cape, South Africa, with the aim of identifying early-life factors that influence child health, development, and illness (42–44). The cohort comprises a low socioeconomic community with a high prevalence of HIV infection and related risk factors. Pregnant women were enrolled between 2012 and 2015 during their second trimester and have been followed, along with their children, from birth through childhood. Eligible women were at least 18 years old, in their second trimester (gestational period of 20-28 weeks), planned to attend one of the two recruitment clinics, and intended to remain in the area. Written informed consent was obtained from all participants and renewed annually.

For this study, serum markers were measured in a subset of mothers during pregnancy (n=138; 60 living with HIV and 78 without HIV) and in their children at 6 weeks (n=97; 41 HEU and 56 HU) and 2 years of age (n=111; 46 HEU and 65 HU). Selection of makers to include in our panel was based on published evidence from pediatric HIV exposure studies reporting alterations in circulating blood markers (**Supplementary Table 1**). A subgroup of children (n=156; 66 HEU and 90 HU) also participated in a longitudinal neuroimaging sub-study at the Cape Universities Body Imaging Centre (CUBIC) (**Supplementary Figure 1**). Participants included in this report were children aged 2–3 years who had undergone neonatal imaging (44), as well as additional children selected based on maternal HIV and alcohol use during pregnancy, and a randomly selected comparison group. Exclusion criteria included medical comorbidities (e.g., congenital abnormalities, genetic syndromes, neurological disorders), low Apgar scores (<7 at 5 minutes), neonatal intensive care admission, maternal use of illicit drugs during pregnancy, child HIV infection, and magnetic resonance imaging (MRI) contraindications such as cochlear implants (45).

### Sociodemographic data collection

Maternal HIV status was determined through routine testing during pregnancy and re-checked every 12 weeks, following the Western Cape PMTCT guidelines at the time (46). HEU children were tested at 6 weeks, 9 months, and 18 months using PCR, rapid antibody, or ELISA tests and confirmed HIV-negative at 18 months or after breastfeeding cessation if this extended beyond then. HU children were defined as those born to mothers without HIV infection. Mothers living with HIV received ART according to PMTCT guidelines, and HEU children received post-exposure prophylaxis from birth (47). Maternal CD4 cell count and viral load data during pregnancy were obtained from clinical records and the National Health Laboratory Service system, using the lowest maternal CD4 cell count within one year before birth and three months after birth to maximize data availability.

Sociodemographic and maternal psychosocial data were collected between 28 and 32 weeks of gestation using interviews and standardized questionnaires adapted from the South African Stress and Health study (42,43). Infant birthweight and nutrition markers were recorded following WHO Z-score guidelines (48). Maternal alcohol use during pregnancy was assessed using the Alcohol, Smoking, and Substance Involvement Screening Test (ASSIST), and retrospectively recorded as a binary measure of moderate-severe alcohol use (44). During pregnancy, maternal smoking was self-reported and assessed by ASSIST, and maternal depression was assessed using the Edinburgh Postnatal Depression Scale, with depression subsequently categorized as a binary variable based on a predefined cutoff.

### Immune assays

Serum samples were collected from mothers at 26–28 weeks of pregnancy and from children at 6 weeks and 2 years of age (42). Concentrations of GM-CSF, INF-γ, IL-1β, IL-2, IL-5, IL-6, IL-7, IL-8, TNF-α, IL-4, IL-10, IL-12p70, and IL-13 were measured using a Milliplex^®^ Luminex 13-plex kit (#HSTCMAG28SPMX13; Merck) according to the manufacturer’s instructions (9). Plates were read on a Luminex system (Bio-Plex 200 System; commercial provider Bio-Rad). ELISA assays (R&D Systems, Minneapolis, USA) were used to measure neutrophil gelatinase-associated lipocalin (NGAL), matrix metalloproteinase-9 (MMP-9), chitinase-3-like kinase (YKL-40), and soluble markers of monocyte activation (sCD14, sCD163) (9). All samples were assayed in duplicate.

Neuroinflammatory markers were selected due to their roles in HIV-related neuropathology. NGAL is involved in neuroinflammation (49) and microglial activation (50) and has been linked to reduced brain volumes and cognitive impairment in a South African cohort of adults living with HIV (30,51). Circulating MMP-9 levels have been found to be significantly lower in adults with HIV compared to controls (51). YKL-40 has been associated with axonal injury (52) and cognitive impairment (53) in cohorts of patients living with HIV from Sweden and the US, respectively.

### Magnetic Resonance Spectroscopy protocol

Participants in the neuroimaging sub-study underwent a multimodal MRI protocol without sedation at the Cape Universities Body Imaging Centre, University of Cape Town, between January 2016 and September 2018. Imaging was performed using a 3 Tesla Siemens Skyra MRI scanner (Erlangen, Germany) with a 32-channel head coil (45). Children were scanned during natural sleep after obtaining informed consent from their parent/legal guardian and ensuring a comfortable and secure positioning using pillows, blankets, and ear protection. A trained study staff member remained in the scanner room throughout the session in case the child awoke.

The MRS acquisition protocol has been previously described (25). Briefly, single-voxel MRS was used to target the midline parietal grey matter as well as bilateral parietal white matter. Spectral quality was maintained using automatic shimming and manual adjustments to reduce spectral linewidths. High-resolution T1-weighted structural images were also acquired for voxel placement.

### Magnetic Resonance Spectroscopy data processing

MRS data were processed using LCModel software (version 6.3-1) for metabolite quantification, as previously described (25,54). Voxels were registered to T1-weighted structural images and segmented into grey matter, white matter, and cerebrospinal fluid to correct for partial volume effects. Metabolite ratios to total creatine were calculated for myo-inositol, glutamate, N-acetyl-aspartate, and total choline.

Spectral quality was visually inspected and assessed using signal-to-noise ratio (SNR) and full width at half maximum (FWHM) given by LCModel. Spectra with FWHM values greater than 0.08 or SNR values lower than 10 were considered of low quality and therefore excluded. Metabolite ratios were compared across brain regions to assess neuroinflammation (myo-inositol), neuronal function (glutamate), neuronal health (n-acetyl-aspartate), and white matter maturation (total choline) (23–25).

### Statistical analysis

Sociodemographic characteristics of the mother-child pairs were reported as mean (±SD) and frequencies (%). Histograms were used to visually inspect the distribution of continuous variables. Differences between HEU and HU children were assessed using independent *t*-tests or Wilcoxon rank-sum tests for normally and non-normally distributed continuous data, respectively. Categorical variables were compared using Chi-square (X^2^) tests.

Serum marker concentrations were log-transformed to meet assumptions of normality. Preliminary group comparisons were performed using independent *t*-tests or Wilcoxon tests, depending on data distribution. Longitudinal trajectories of child peripheral blood markers from 6 weeks to 2 years of age were analyzed via LMMs with Maximum Likelihood Estimation with use of the *lme4* R package. This approach allowed us to account for intra-individual variability and repeated measures, without requiring complete cases or imputation, making it suitable for handling missing data across timepoints. Three covariance structures —variance components, compound symmetry, and unstructured— were considered to determine the best model fit using AIC and BIC values.

Linear regression models with robust standard errors and BH correction for multiple comparisons were applied to investigate cross-sectional associations between serum markers and brain metabolite ratios in HEU and HU children at age 2–3 years. For MRS data, a factor analysis approach was used to construct weighted linear combinations of all four metabolite ratios across three parietal brain voxels, creating four metabolite patterns —composite factor scores—, as previously described (25). Separate models were then run to investigate cross-sectional associations between serum and metabolite patterns. Adjusted linear models were run for associations that remained statistically significant after BH correction. Potential confounders were chosen a priori due to their reported influence in neurometabolite or neurobehavioral outcomes in children, and included child age (23,24), and child sex (55,56). Given that the percentage of white matter in white matter voxels was suboptimal in our study (≈52), tissue composition was also included in the adjusted analysis (25). Sensitivity analyses were conducted for any sociodemographic or clinical variables that showed significant differences between HEU and HU groups.

Mediation analyses were performed on serum markers and neurometabolites with statistically significant associations in the adjusted linear models, using the *lavaan* R package for structural equation modelling. Bootstrapping was used to compute confidence intervals for the indirect effect, providing robust estimates that did not rely on the assumption of a normal distribution. This approach was particularly valuable in our sample, where small subsample sizes limit statistical power.

Results were considered significant at a p-value threshold of < 0.05 (two-tailed). Statistical analyses were conducted using R (version 4.4.2) with RStudio software. The analysis plan was pre-registered in the Open Science Framework (57).

## Ethics approval and consent

The studies involving human participants were reviewed and approved by the Faculty of Health Sciences, Human Research Ethics Committee, University of Cape Town (401/2009; 525/2012 & 044/2017), by Stellenbosch University (N12/02/0002), and by the Western Cape Provincial Health Research committee (2011RP45). Written informed consent to participate in this study was provided by the participants’ parent/legal guardian.

## Author Contributions

**CBC**: methodology, formal analysis and interpretation, visualization, writing – original draft, review & editing. **FR**: methodology, formal analysis, supervision, and writing – review & editing. **TK**: methodology, data curation and writing – review & editing. **JA**: methodology, data curation and writing – review & editing. **SS**: investigation and writing – review & editing. **KN**: methodology and writing – review & editing. **SJ**: methodology and writing – review & editing. **NH**: project administration and writing – review & editing. **HZ**: conceptualization, methodology, resources, and writing - review & editing. **DS**: conceptualization, methodology, investigation, resources, supervision, and writing – review & editing. **KD**: conceptualization, methodology, investigation, resources, supervision, and writing – review & editing. **CW**: conceptualization, methodology, investigation, data curation, supervision, and writing – review & editing. **PN**: conceptualization, methodology, investigation, data curation, supervision, and writing – review & editing.

All authors approved the final version.

## Data Availability

The Drakenstein Child Health Study is committed to the principle of data sharing. De-identified data will be made available to requesting researchers as appropriate. Requests for collaborations to undertake data analysis are welcome.

More information can be found on our website http://www.paediatrics.uct.ac.za/scah/dclhs. A comprehensive Statistical Analysis Report, as well as the R code used for this study analysis, are openly accessible on GitHub: https://github.com/Cescualito/UCT_DCHS_PhD.

## Acknowledgments

In memory of Dr Annerine Roos. We show our deep gratitude to the mothers and their children for participating in the Drakenstein Child Health Study. We thank the research nurses Tabitha Mutseyekwa and Judy Gatei, and research assistant Joavine Fourie for their work, the clinical and administrative staff of the Western Cape Government Health Department at Paarl Hospital and at the clinics for support of the study. We are also grateful for the assistance provided by the Cape Universities Body Imaging Centre (CUBIC) team, particularly Petty Samuels, Ingrid Op’t Hof and Mazwi Maishi.

