## Supplementary materials for "Maternal and child immune profiles are associated with neurometabolite measures of early-life neuroinflammation in children who are HIV-exposed and uninfected: a South African birth cohort"

[‡]Shared last authorship

#### Supplementary Table 1. Studies reporting peripheral blood immune marker alterations in mothers living with HIV and/or children who are HIV-exposed and uninfected

**Legend**: Children (a); Pregnant women (b)

**Controls** are either defined as (a) children born to mothers who are not living with HIV; or (b) pregnant women not living with HIV

**Cases** are either defined as (a) children born to mothers who are living with HIV; or (b) pregnant women living with HIV

| **Study**, **year** | | **Country** | **Technique** | **N controls** | **N cases** | **Timepoint / Age** | **Reported differences in serum marker levels between groups** |
|---|---|---|---|---|---|---|---|
| (1) | Sachdeva *et al.,* 2008 | United States | Biochip Array | 15 | 35 | Pregnancy, gestational age 12–15 weeks | **TNFα** was higher in women living with HIV compared to women without HIV. |
| (2) | Richardson *et al*., 2011 | United States | ELISA & CLIA | 18 | 20 | Pregnancy, 2nd and 3rd trimesters | **IFN-γ**, **IL1**, **IL4**, **IL8**, **IL10**, and **TNFα** were higher in women living with HIV compared to women without HIV. There were no significant changes in plasma cytokines and other biomarkers from early to late pregnancy. |
| (3) | López *et al*., 2016 | Spain | ELISA | 36 | 36 | 1st trimester | In both trimesters, **CD14** was higher in women living with HIV compared to women without HIV. |
| | | | | | | 3rd trimester | |
| (4) | Maharaj *et al.,* 2017 | South Africa | CBA | 50 | 45 | Pregnancy, gestational age 35–36 weeks | **IL-2**, **IL-6**, and **TNF-α** were lower in women living with HIV compared to women without HIV. |
| (5) | Prendergast *et al*., 2017 | Zimbabwe | ELISA | 197 | 194 | 6 weeks | No group differences in **IL-6** levels were detected at any timepoint. |
| | | | | | | 6 months | |
| (6) | Evans *et al*., 2017 | Zimbabwe | ELISA | 97 | 223 | 6 weeks | **C-reactive protein** was significantly higher in HEU infants compared to their HU peers. |
| (7) | Miyamoto *et al*., 2017 | Brazil | Luminex & ELISA | 20 | 19 | Birth | No group differences were detected. |
| | | | | 19 | 19 | 12 months | A slight decay in **CD14** levels was observed from 12 months to 6–12 years in HEU children. At 6–12 years, **IL-4** was higher in HEU compared to HU children. |
| | | | | 18 | 20 | 6–12 years | |
| (8) | Dirajlal-Fargo *et al.*, 2019 | Brazil | ELISA | 88 | 86 | Delivery | **IL-6** and **CD14** were higher in women living with HIV compared to women without HIV. |
| | | | | 88 | 86 | Birth | **IL-6** and **CD14** were higher in HEU compared to HU infants. |
| | | | | | | 6 months | **IL-6** remained significantly higher in HEU infants. |
| (9) | Ray *et al*., 2019 | Kenya | Luminex | 43 | 44 | Delivery | **IFN-γ**, **IL-1β**, **IL-6**, **IL-10**, **IL-12p70**, **IL-17A**, **IL-17E**, **IL-17F**, **IL-21**, **IL-22**, **IL-23**, and **TNFα** were significantly lower in mothers with HIV compared to those without HIV. |

| | | | | | | | |
|---|---|---|---|---|---|---|---|
| | | | | 43 | 44 | Birth | No difference was detected for any cytokine measured in cord blood between HEU and HU neonates. |
| (10) | Shafiq *et al*., 2021 | India | Luminex | 149 | 69 | Pregnancy, gestational age 28–30 weeks | Higher **IL-1β** levels during pregnancy were associated with preterm birth in women living with and without HIV. Higher **CD14** levels during pregnancy were associated with growth deficits at birth in mothers living with HIV. |
| (11) | Sevenoaks *et al*., 2021 | South Africa | Luminex & ELISA | 190 | 77 | Pregnancy, gestational age ≈26 weeks | **GM-CSF** and **MMP-9** were lower in mothers living with HIV compared to mothers without HIV. **IL-1β** and **IL-4** were also lower in mothers living with HIV prior to correction for multiple comparisons.<br>In mothers living with HIV, and prior to correction for multiple comparisons, **IFN-γ**, **IL-10**, **IL-12p70** and **IL-7** were associated with lower composite scores for language in HEU children at 24–28 months, and **TNF-α** was associated with lower cognitive scores. |
| | | | | 159 | 63 | 6–10 weeks | **IFN-γ** and **IL-1β** were lower in HEU compared to HU infants. **IL-12p70** and **IL-4** were also lower in HEU infants prior to correction for multiple comparisons.<br>In HEU infants, **GM-CSF**, **IFN-γ**, **IL-10**, **IL-12p70**, **IL-1β**, **IL-2**, **IL-4**, **IL-6**, and **NGAL** were associated with motor development at 24–28 months. Prior to correction for multiple comparisons, **MMP-9** was associated with motor and language outcomes, and **IL-1β** was associated with language outcomes. |
| | | | | 190 | 77 | 24–28 months | **IFN-γ**, **IL-1β**, **IL-2** and **IL-4** were lower in HEU compared to HU children. In HEU children, and prior to correction for multiple comparisons, **IL-10** was associated with lower cognitive scores. |
| (12) | Akoto *et al*., 2021 | South Africa | Luminex | 68 | 56 | Pregnancy, 1st trimester | Detection of **IL-1β**, **IFN-β** and **IFN-λ2/3** was lower in women living with HIV compared to those without HIV. Detection of **IP-10**, **IL-2**, **IL-5**, **IL-6**, **IL-9**, **IL-10**, and **IL-17A** was higher in women living with HIV compared to those without HIV. |
| | | | | | | 2nd trimester | Detection of **IL-1β**, **IFN-β** and **IFN-λ2/3** was lower in women living with HIV compared to those without HIV. Detection of **IFN-λ1**, **IP-10**, **IL-2**, **IL-5**, **IL-10**, **IL-12p70**, and **IL-17A** was higher in women living with HIV compared to those without HIV. |
| | | | | | | 3rd trimester | Detection of **IFN-β** and **IFN-λ2/3** was lower in women living with HIV compared to those without HIV. Detection of **IFN-λ1**, **IP-10**, and **IL-5** was higher in women living with HIV compared to those without HIV. |
| (13) | Schnittman *et al*., 2021 | Uganda | ELISA | — — | 759 | Pre-pregnancy, 1st, 2nd, and 3rd trimester pregnancy, and postpartum | **IL-6** declined by 29% in the 1st trimester but increased toward pre-pregnancy baseline by the 3rd trimester. **CD14** declined by 17%–18% in the 1st and 2nd trimesters. **CD27** and **CD163** declined in the 1st trimester. **IP-10** declined by 30%–40% during pregnancy. |
| (14) | Vyas *et al*., 2021 | India | Luminex | 150 | 70 | Pregnancy, gestational age 13–34 weeks | In the second trimester, **CD14**, **TNFα**, **IL-6**, and **IL-17a** were higher, and **CD163** lower, in pregnant women living with HIV compared to women without HIV. In the third trimester, **CD14** and **IL-6** were higher in women living with HIV compared to women without HIV. |

| | | | | | | | |
|---|---|---|---|---|---|---|---|
| (15) | Shiau *et al*., 2023 | United States | EIA | 76 | 188 | Pregnancy, gestational age 13–27 weeks | **IL-6**, **CD14**, and **CD163** were higher in pregnant people living with HIV compared to those without HIV. Among people living with HIV, **CD14** and **CD163** were higher in those with perinatally acquired HIV versus non-perinatally acquired HIV. |
| (16) | Bebell *et al*., 2024 | Uganda | Luminex | 142 | 147 | Delivery | Partial Least Squares Discriminant Analysis identified top markers distinguishing cytokine profiles, which included higher **IL-5** in pregnant women living with HIV, higher **IL-8** and **MIP-1α** in women without HIV, and higher **RANTES** and **E-selectin** in umbilical cord plasma from HU newborns. |
| (17) | Ray *et al*., 2024 | Kenya | Luminex | 58 | 59 | From birth to 54 weeks of age | There were no significant interaction effects between maternal HIV status and time for any of the cytokines measured, indicating that cytokine trajectories did not differ between HEU and HU children. |
| (18) | Hindle *et al*., 2024 | Canada | Luminex | 22 | 144 | Pregnancy, 2nd and 3rd trimesters | In both trimesters, **AGP** was higher and **IFN-β** lower in pregnant people living with HIV compared to those without HIV. In the second trimester, **HMGB1**, **IFN-γ**, and **IFN-α** were lower in pregnant people living with HIV compared to those without HIV. |
| (19) | Yin *et al*., 2024 | United States | Luminex | 18 | 46 | Within 2 days prior to delivery | **IL-1β**, **IL-21**, **TNF-α**, **CCL5**, **CXCL9**, **sCD27**, **sCD40L**, and **sCD163** were higher in pregnant women living with HIV compared to those without HIV, whereas **APRIL** was lower. **CXCL9** and **CXCL10** were significantly higher in pregnant women living with HIV who were not virally suppressed compared to those who were virally suppressed. |
| | | | | 50 | 46 | Birth | **IL-1β**, **IL-6**, **TNF-α**, **IL-10**, **IL-1RA**, **IL-21**, **IL-22**, **CCL4**, **CXCL9**, **sCD14**, **sCD27**, **sCD40L**, **sCD163**, and **APRIL** were significantly higher in HEU newborns compared to HU. |
| | | | | 50 | 46 | 6 months | **IL-1β**, **TNF-α**, **IL-21**, **CCL4**, **sCD14**, **sCD40L**, and **APRIL** were significantly higher in HEU children compared to HU. |

**CLIA**: Chemiluminescence Immunoassay; **ELISA**: Enzyme-Linked Immunosorbent Assay; **CBA**: Cytometric Bead Array; **EIA**: Enzyme Immunoassay; **HEU**: HIV-exposed uninfected; **HU**: HIV-unexposed.

#### 2 Supplementary Table 2

##### 2A. Sociodemographic characteristics of the subset of children included in the MRS analysis

| | **DCHS complete MRS subset** (**N=83**) | | |
|---|---|---|---|
| | **CHEU** (**n=36**) Median (IQR) or n/N (%) | **CHU** (**n=47**) Median (IQR) or n/N (%) | **P value** |
| **Child age at scan** (in months) | 34.00 (2.00) | 35.00 (1.00) | 0.21 |
| **Sex** | | | 0.14 |
| Female | 11/36 (30.6%) | 23/47 (48.9%) | |
| Male | 25/36 (69.4%) | 24/47 (51.1%) | |
| **Monthly household income** (in ZAR) | | | 0.98 |
| <R1,000 | 10/36 (27.8%) | 14/47 (29.8%) | |
| R1,000–5,000 | 23/36 (63.9%) | 29/47 (61.7%) | |
| >R5,000 | 3/36 (8.3%) | 4/47 (8.5%) | |
| **Maternal education** | | | 0.83 |
| Primary | 3/36 (8.3%) | 3/47 (6.4%) | |
| Some secondary | 22/36 (61.1%) | 26/47 (55.3%) | |
| Completed secondary | 10/36 (27.8%) | 15/47 (31.9%) | |
| Tertiary | 1/36 (2.8%) | 3/47 (6.4%) | |
| **Employed mother** | 9/36 (25.0%) | 9/47 (19.1%) | 0.71 |
| **Maternal relationship status** (partnered) | 19/36 (52.8%) | 17/47 (36.2%) | 0.20 |
| **Maternal age at delivery** (in years) | 29.49 (6.15) | 24.93 (6.74) | **<0.0001** |
| **Gestational age at delivery** (in weeks) | 39.00 (2.25) | 39.00 (2.00) | 0.66 |
| **Premature birth** (<37 weeks' gestation) | 5/36 (13.9%) | 6/47 (12.8%) | 1.00 |
| **Child birthweight** (in grams) | 3120.00 (547.50) | 3210.00 (645.00) | 0.26 |
| **Exclusive breastfeeding** for 5 or more months | 7/36 (19.4%) | 4/47 (8.5%) | 0.26 |
| **Exclusive breastfeeding duration** (in months) | 0.92 (3.10) | 1.84 (2.04) | 0.07 |
| **Nutritional status at 2 years old** | | | |
| Stunting (height-for-age Z-score < -2) | 6/31 (19.4%) | 3/43 (7.0%) | 0.20 |
| Underweight (weight-for-age Z-score < -2) | 1/32 (3.1%) | 1/43 (2.3%) | 0.90 |
| Wasting (weight-for-length Z-score < -2) | 1/32 (3.1%) | 1/43 (2.3%) | 0.90 |

| | | | |
|---|---|---|---|
| **Maternal anaemia during pregnancy** | 11/36 (30.6%) | 14/47 (29.8%) | 1.00 |
| **Maternal smoking during pregnancy** | 7/36 (19.4%) | 17/47 (36.2%) | 0.16 |
| **Maternal alcohol use during pregnancy** | 4/34 (11.8%) | 11/46 (23.9%) | 0.28 |
| **Maternal depression during pregnancy** | 1/28 (3.6%) | 11/42 (26.2%) | **0.019** |
| **Maternal hospitalization during pregnancy** | 3/36 (8.3%) | 4/47 (8.5%) | 1.00 |
| **Maternal HIV diagnosis timepoint** | | | |
| Before pregnancy | 16/36 (44.4%) | | |
| During pregnancy | 20/36 (55.6%) | | |
| **Maternal lowest CD4 cell count during pregnancy**[§] | | | |
| ≤500 cells/mm$^3$ | 13/26 (50.0%) | | |
| >500 cells/mm$^3$ | 13/26 (50.0%) | | |
| **Highest maternal viral load during pregnancy** | | | |
| (undetectable) <40 copies/mL | 24/29 (82.8%) | | |
| 40–1000 copies/mL | 2/29 (6.9%) | | |
| >1000 copies/mL | 3/29 (10.3%) | | |
| **Antiretroviral therapy initiation** | | | |
| Before pregnancy | 16/36 (44.4%) | | |
| During pregnancy | 20/36 (55.6%) | | |
| **First-line antiretroviral therapy during pregnancy** | | | |
| Efavirenz + Emtricitabine + Tenofovir (FDC) | 33/36 (91.7%) | | |
| Lamivudine + Zidovudine + Nevirapine | 2/36 (5.6%) | | |
| Lamivudine + Zidovudine + Efavirenz | 1/36 (2.8%) | | |
| **Cotrimoxazole prophylaxis** | 31/32 (96.9%) | | |
| **Infant prophylaxis** | | | |
| Nevirapine monotherapy | 28/36 (77.8%) | | |
| Nevirapine + zidovudine | 8/36 (22.2%) | | |

Data are median (IQR) or n/N (%). Percentages calculated out of available data. Continuous data was assessed for normality using Shapiro-Wilk tests. Comparisons between CHEU and CHU were made using Wilcoxon Rank Sum (Mann Whitney U) tests for continuous data, and $X^2$ tests for categorical data. **DCHS**: Drakenstein Child Health Study; **CHEU**: Children who are HIV-Exposed and Uninfected; **CHU**: Children who are HIV-Unexposed; **ZAR**: South African Rand; **FDC**: Fixed Dose Combination.

Missing data: nutritional conditions at 2 years old (n=5 in the CHEU group, n=4 in the CHU group); maternal alcohol use during pregnancy (n=2 in the CHEU group, n=1 in the HU group); maternal depression during pregnancy (n=8 in the CHEU group, n=5 in the HU group); maternal lower CD4 cell count during pregnancy (n=10); maternal highest viral load during pregnancy (n=7); cotrimoxazole prophylaxis (n=4). [§]The lowest maternal CD4 cell count within 1 year before birth and 3 months after birth was used to maximise numbers.

#### 2B. Sociodemographic characteristics of the subset of children included in the MRS analysis, compared to all children invited for neuroimaging at age 2–3 years

| | **DCHS neuroimaging cohort at age 2 years** | | |
|---|---|---|---|
| | **Complete MRS subset (N=83)** Median (IQR) or n/N (%) | **Original cohort (N=156)** Median (IQR) or n/N (%) | **p value** |
| **Child age at scan** (in months) | 34.00 (2.00) | 34.00 (2.00) | 0.71 |
| **Sex** | | | 0.95 |
| Male | 49/83 (59.0%) | 90/156 (57.7%) | |
| Female | 34/83 (41.0%) | 66/156 (42.3%) | |
| **Monthly household income** (in ZAR) | | | 0.67 |
| <R1,000 | 24/83 (28.9%) | 37/156 (23.7%) | |
| R1,000–5,000 | 52/83 (62.7%) | 104/156 (66.7%) | |
| >R5,000 | 7/83 (8.4%) | 15/156 (9.6%) | |
| **Maternal education** | | | 0.96 |
| Primary | 6/83 (7.2%) | 9/156 (5.8%) | |
| Some secondary | 48/83 (57.8%) | 92/156 (59.0%) | |
| Completed secondary | 25/83 (30.1%) | 46/156 (29.5%) | |
| Tertiary | 4/83 (4.8%) | 9/156 (5.8%) | |
| **Employed mother** | 18/83 (21.7%) | 42/156 (26.9%) | 0.46 |
| **Maternal relationship status** (partnered) | 36/83 (43.4%) | 74/156 (47.4%) | 0.64 |
| **Maternal age at delivery** (in years) | 27.09 (7.43) | 27.26 (7.56) | 0.69 |
| **Gestational age at delivery** (in weeks) | 39.00 (2.00) | 39.00 (2.00) | 0.99 |
| **Premature birth** (<37 weeks' gestation) | 11/83 (13.3%) | 20/156 (12.8%) | 1.00 |
| **Birthweight** (in grams) | 3170.00 (540.00) | 3180.00 (657.50) | 0.87 |
| **Exclusive breastfeeding** for 5 or more months | 11/83 (13.3%) | 18/156 (11.5%) | 0.86 |
| **Exclusive breastfeeding duration** (in months) | 1.84 (2.42) | 1.00 (2.54) | 0.31 |
| **Nutritional status at 2 years old** | | | |
| Stunting (height-for-age Z-score < -2) | 9/74 (12.2%) | 18/138 (13.0%) | 0.97 |
| Underweight (weight-for-age Z-score < -2) | 2/75 (2.7%) | 134/139 (96.4%) | 0.89 |
| Wasting (weight-for-length Z-score < -2) | 2/75 (2.7%) | 5/139 (3.6%) | 0.89 |

| | | | |
|---|---|---|---|
| **Maternal anaemia during pregnancy** | 25/83 (30.1%) | 48/156 (30.8%) | 1.00 |
| **Maternal smoking during pregnancy** | 24/83 (28.9%) | 41/156 (26.3%) | 0.78 |
| **Maternal alcohol use during pregnancy** | 15/80 (18.8%) | 25/152 (16.4%) | 0.82 |
| **Maternal depression during pregnancy** | 12/70 (17.1%) | 30/128 (23.4%) | 0.53 |
| **Maternal hospitalization during pregnancy** | 7/83 (8.4%) | 10/154 (6.5%) | 0.50 |
| **Maternal HIV status** | | | 0.98 |
| Positive | 36/83 (43.4%) | 66/156 (42.3%) | |
| Negative | 47/83 (56.6%) | 90/156 (57.7%) | |

Data are median (IQR) or n/N (%). Percentages calculated out of available data. Continuous data was assessed for normality using Shapiro-Wilk tests. Comparisons between CHEU and CHU were made using Wilcoxon Rank Sum (Mann Whitney U) for continuous data, and $X^2$ tests for categorical data. **DCHS**: Drakenstein Child Health Study; **CHEU**: Children who are HIV-Exposed and Uninfected; **CHU**: Children who are HIV-Unexposed; **ZAR**: South African Rand; **FDC**: Fixed Dose Combination.

Missing data: nutritional conditions at 2 years old (n=9 in the complete MRS subset, n=18 in the full cohort); maternal hospitalization during pregnancy (n=2 in the full cohort); maternal alcohol use during pregnancy (n=3 in the complete MRS subset, n=4 in the full cohort); maternal depression during pregnancy (n=13 in the complete-case cohort, n=28 in the original cohort).

### 3 Supplementary Table 3. Maternal, infant, and child serum marker concentrations

#### 3.1 Maternal serum marker concentrations during pregnancy (log-scaled)

| Biomarker | Mothers not living with HIV (n=78) | Mothers living with HIV (n=60) | Effect size | 95% CI | | P-value | BH |
|---|---|---|---|---|---|---|---|
| GM-CSF | 3.76 ± 0.98 | 3.29 ± 0.89 | 0.47 | 0.16 | 0.79 | **0.004** | **0.034** |
| IFN-γ | 2.27 (1.1) | 2.29 (1.45) | -0.08 | -0.33 | 0.18 | 0.54 | 0.88 |
| IL-1β | 0.58 (1.02) | 0.54 (1.32) | 0.09 | -0.18 | 0.36 | 0.51 | 0.88 |
| IL-2 | 0.77 (1.18) | 0.78 (1.58) | 0.02 | -0.30 | 0.34 | 0.93 | 0.98 |
| IL-5 | 0.76 ± 0.77 | 0.82 ± 0.88 | -0.06 | -0.35 | 0.22 | 0.66 | 0.90 |
| IL-6 | 0.67 (1.59) | 0.50 (1.64) | 0.00 | -0.45 | 0.44 | 0.98 | 0.98 |
| IL-7 | 2.31 (0.8) | 2.25 (0.92) | -0.01 | -0.21 | 0.19 | 0.88 | 0.98 |
| IL-8 | 1.28 (0.91) | 1.26 (0.88) | -0.06 | -0.32 | 0.20 | 0.67 | 0.98 |
| TNFα | 1.58 ± 0.55 | 1.71 ± 0.57 | -0.14 | -0.33 | 0.06 | 0.16 | 0.49 |
| IL-4 | 3.51 (1.68) | 2.99 (2.06) | 0.32 | -0.13 | 0.73 | 0.18 | 0.66 |
| IL-10 | 2.18 ± 1.01 | 2.15 ± 1.08 | 0.03 | -0.32 | 0.39 | 0.85 | 0.90 |
| IL-12p70 | 1.25 (1.28) | 1.24 (1.18) | 0.03 | -0.23 | 0.32 | 0.77 | 0.98 |
| IL-13 | 1.55 (1.43) | 1.19 (1.64) | 0.25 | -0.17 | 0.67 | 0.26 | 0.77 |
| CD14 | 7.46 ± 0.34 | 7.59 ± 0.38 | -0.13 | -0.25 | 0.00 | **0.044** | 0.20 |
| CD163 | 6.32 ± 0.48 | 6.31 ± 0.5 | 0.02 | -0.15 | 0.19 | 0.82 | 0.90 |
| NGAL | 5.21 ± 0.51 | 4.99 ± 0.56 | 0.22 | 0.04 | 0.40 | **0.018** | 0.11 |
| MMP-9 | 7.11 ± 0.62 | 6.64 ± 0.75 | 0.48 | 0.24 | 0.72 | **0.0001** | **0.002** |
| YKL-40 | 3.51 (0.87) | 3.61 (1.03) | -0.12 | -0.38 | 0.13 | 0.32 | 0.77 |

*T*-Test for normally-distributed data; Wilcoxon Rank-Sum Test (Mann-Whitney U Test) for not normally-distributed data. Data presented as mean ±SD or median (IQR) per each group.

**BH**: Benjamini-Hochberg corrected p-value.

#### 3.2 Infant serum marker concentrations at 6 weeks of age (log-scaled)

| Biomarker | CHU (n=56) | CHEU (n=41) | Effect size | 95% CI | | P-value | BH |
|---|---|---|---|---|---|---|---|
| GM-CSF | 2.68 (1.07) | 2.35 (1.08) | 0.24 | -0.11 | 0.59 | 0.19 | 0.86 |
| IFN-γ | 1.43 ± 0.97 | 1.41 ± 1.03 | 0.02 | -0.39 | 0.44 | 0.91 | 0.99 |
| IL-1β | -0.08 ± 1.05 | -0.42 ± 1.00 | 0.34 | -0.08 | 0.76 | 0.11 | 0.66 |
| IL-2 | 0.01 (1.46) | 0.10 (1.43) | 0.01 | -0.41 | 0.45 | 0.95 | 0.97 |
| IL-5 | 0.41 (0.96) | 0.44 (0.89) | -0.12 | -0.47 | 0.20 | 0.42 | 0.97 |
| IL-6 | 0.49 (2.08) | 0.35 (1.84) | 0.01 | -0.55 | 0.56 | 0.96 | 0.97 |
| IL-7 | 1.68 (0.96) | 1.76 (0.82) | -0.09 | -0.36 | 0.16 | 0.49 | 0.97 |
| IL-8 | 1.86 (0.65) | 2.01 (1.07) | -0.08 | -0.35 | 0.21 | 0.56 | 0.97 |
| TNFα | 2.79 ± 0.42 | 2.87 ± 0.71 | -0.08 | -0.33 | 0.17 | 0.52 | 0.99 |
| IL-4 | 2.20 (2.62) | 1.93 (2.02) | 0.08 | -0.47 | 0.64 | 0.77 | 0.97 |
| IL-10 | 2.46 ± 0.82 | 2.49 ± 0.65 | -0.03 | -0.32 | 0.27 | 0.86 | 0.99 |
| IL-12p70 | 0.60 ± 1.00 | 0.45 ± 0.85 | 0.15 | -0.22 | 0.52 | 0.43 | 0.99 |
| IL-13 | 1.24 (1.70) | 1.10 (1.50) | 0.04 | -0.45 | 0.57 | 0.88 | 0.97 |
| CD14 | 7.38 ± 0.28 | 7.46 ± 0.26 | -0.09 | -0.20 | 0.02 | 0.11 | 0.66 |
| CD163 | 6.41 ± 0.45 | 6.46 ± 0.50 | -0.05 | -0.25 | 0.14 | 0.60 | 0.99 |
| NGAL | 4.50 (0.53) | 4.32 (0.47) | 0.19 | 0.02 | 0.35 | **0.032** | 0.51 |
| MMP-9 | 5.79 (0.51) | 5.71 (1.05) | 0.07 | -0.22 | 0.35 | 0.66 | 0.97 |
| YKL-40 | 3.43 ± 0.53 | 3.37 ± 0.46 | 0.06 | -0.14 | 0.26 | 0.56 | 0.99 |

*T*-Test for normally-distributed data; Wilcoxon Rank-Sum Test (Mann-Whitney U Test) for not normally-distributed data. Data presented as mean ±SD or median (IQR) per each group.

**CHU**: Children who are HIV-Unexposed; **CHEU**: Children who are HIV-Exposed and Uninfected; **BH**: Benjamini-Hochberg corrected p-value.

#### 3.3 Child serum marker concentrations at 2 years of age (log-scaled)

| Biomarker | CHU (n=65) | CHEU (n=46) | Effect size | 95% CI | | P-value | BH |
|---|---|---|---|---|---|---|---|
| GM-CSF | 4.62 ± 0.88 | 4.44 ± 0.85 | 0.18 | -0.15 | 0.52 | 0.27 | 0.63 |
| IFN-γ | 2.12 ± 0.68 | 1.97 ± 0.65 | 0.15 | -0.10 | 0.41 | 0.24 | 0.63 |
| IL-1β | 0.71 (1.57) | 0.13 (1.13) | 0.33 | 0.04 | 0.66 | **0.031** | 0.27 |
| IL-2 | 0.89 (1.30) | 0.63 (0.87) | 0.28 | -0.01 | 0.55 | 0.06 | 0.27 |
| IL-5 | 1.06 (0.85) | 1.06 (0.71) | 0.03 | -0.19 | 0.27 | 0.75 | 0.82 |
| IL-6 | 1.20 (0.83) | 0.95 (0.96) | 0.17 | -0.16 | 0.46 | 0.28 | 0.47 |
| IL-7 | 2.16 ± 0.54 | 2.09 ± 0.43 | 0.07 | -0.11 | 0.26 | 0.43 | 0.65 |
| IL-8 | 2.23 (1.05) | 2.23 (1.43) | 0.08 | -0.31 | 0.44 | 0.62 | 0.75 |
| TNFα | 2.49 (0.72) | 2.46 (0.64) | -0.03 | -0.22 | 0.18 | 0.79 | 0.82 |
| IL-4 | 3.66 (1.18) | 3.28 (1.65) | 0.36 | -0.01 | 0.74 | 0.06 | 0.27 |
| IL-10 | 2.82 ± 0.77 | 2.70 ± 0.53 | 0.12 | -0.13 | 0.36 | 0.34 | 0.63 |
| IL-12p70 | 1.55 (1.35) | 1.26 (1.05) | 0.22 | 0.00 | 0.44 | **0.048** | 0.27 |
| IL-13 | 2.23 ± 1.03 | 1.96 ± 0.93 | 0.27 | -0.10 | 0.64 | 0.15 | 0.63 |
| CD14 | 7.71 ± 0.34 | 7.65 ± 0.34 | 0.06 | -0.07 | 0.19 | 0.35 | 0.63 |
| CD163 | 6.52 ± 0.42 | 6.61 ± 0.47 | -0.09 | -0.26 | 0.08 | 0.29 | 0.63 |
| NGAL | 5.21 ± 0.63 | 5.14 ± 0.62 | 0.07 | -0.17 | 0.31 | 0.56 | 0.72 |
| MMP-9 | 6.64 ± 0.64 | 6.69 ± 0.60 | -0.05 | -0.28 | 0.19 | 0.70 | 0.79 |
| YKL-40 | 3.27 (0.83) | 3.36 (0.92) | -0.07 | -0.33 | 0.16 | 0.58 | 0.75 |

*T*-Test for normally-distributed data; Wilcoxon Rank-Sum Test (Mann-Whitney U Test) for not normally-distributed data. Data presented as mean ±SD or median (IQR) per each group.

**CHU**: Children who are HIV-Unexposed; **CHEU**: Children who are HIV-Exposed and Uninfected; **BH**: Benjamini-Hochberg corrected p-value.

**Supplementary Table 4. Linear Mixed-Effects Models to examine child trajectories in serum marker concentrations from 6 weeks to 2 years of age**

| | **Fixed effects** | | | | | | | **Random effects** | | **Model fit** | | | |
|---|---|---|---|---|---|---|---|---|---|---|---|---|---|
| **Serum marker** | **Baseline** log(levels) | **Effect of time** (p-value) | | **Effect of maternal HIV** (p-value) | | **Interaction** (p-value) | | **Intercept** | **Residual SD** | **AIC** | **BIC** | **LogLik** | **Deviance** |
| **GM-CSF** | 0.09 | -0.01 | (0.91) | -0.21 | (0.15) | 0.01 | (0.95) | 0.18 | 0.42 | 594.96 | 614.99 | -291.48 | 582.96 |
| **IFN-γ** | 0.10 | ≈0.00 | (0.95) | -0.08 | (0.63) | -0.12 | (0.42) | 0.72 | 0.69 | 576.56 | 596.58 | -282.28 | 564.56 |
| **IL-1β** | 0.17 | -0.05 | (0.64) | -0.30 | (0.06) | 0.02 | (0.92) | 0.67 | 0.71 | 573.70 | 593.72 | -280.85 | 561.70 |
| **IL-2** | 0.08 | 0.05 | (0.64) | -0.17 | (0.28) | -0.20 | (0.25) | 0.53 | 0.84 | 590.91 | 610.94 | -289.46 | 578.91 |
| **IL-5** | -0.03 | 0.11 | (0.39) | 0.06 | (0.68) | -0.26 | (0.17) | 0.28 | 0.95 | 597.85 | 617.88 | -292.93 | 585.85 |
| **IL-6** | ≈0.00 | -0.02 | (0.90) | ≈0.00 | (0.99) | 0.02 | (0.92) | 0.39 | 0.92 | 599.02 | 619.05 | -293.51 | 587.02 |
| **IL-7** | 0.02 | 0.07 | (0.52) | ≈0.00 | (0.99) | -0.20 | (0.26) | 0.51 | 0.85 | 592.65 | 612.68 | -290.33 | 580.65 |
| **IL-8** | -0.01 | 0.02 | (0.88) | 0.03 | (0.83) | -0.07 | (0.66) | 0.55 | 0.83 | 591.83 | 611.85 | -289.91 | 579.83 |
| **TNFα** | -0.04 | 0.02 | (0.89) | 0.10 | (0.53) | -0.05 | (0.77) | 0.57 | 0.81 | 589.90 | 609.92 | -288.95 | 577.90 |
| **IL-4** | 0.06 | -0.04 | (0.62) | -0.07 | (0.69) | -0.11 | (0.44) | 0.76 | 0.65 | 570.25 | 590.27 | -279.12 | 558.25 |
| **IL-10** | 0.03 | 0.05 | (0.63) | -0.07 | (0.68) | -0.17 | (0.28) | 0.63 | 0.76 | 581.87 | 601.90 | -284.94 | 569.87 |
| **IL-12p70** | 0.07 | 0.05 | (0.89) | -0.13 | (0.26) | -0.16 | (0.66) | 0.63 | 0.76 | 581.21 | 601.23 | -284.60 | 569.21 |
| **IL-13** | 0.07 | 0.03 | (0.79) | -0.16 | (0.29) | -0.13 | (0.48) | 0.52 | 0.84 | 592.29 | 612.32 | -290.15 | 580.29 |
| **CD14** | -0.03 | 0.16 | (0.21) | 0.07 | (0.61) | -0.37 | (**0.06**) | 0.25 | 0.95 | 599.28 | 619.33 | -293.64 | 587.28 |
| **CD163** | -0.08 | -0.02 | (0.88) | 0.15 | (0.34) | 0.08 | (0.63) | 0.55 | 0.83 | 593.15 | 613.20 | -290.57 | 581.15 |
| **NGAL** | 0.08 | -0.07 | (0.53) | -0.24 | (0.11) | 0.16 | (0.34) | 0.48 | 0.86 | 594.53 | 614.58 | -291.26 | 582.53 |
| **MMP-9** | ≈0.00 | -0.05 | (0.70) | ≈0.00 | (0.98) | 0.11 | (0.56) | 0.22 | 0.97 | 602.52 | 622.57 | -295.26 | 590.52 |
| **YKL-40** | ≈0.00 | -0.11 | (0.27) | 0.03 | (0.88) | 0.25 | (0.11) | 0.69 | 0.72 | 581.36 | 601.42 | -284.68 | 569.36 |

**SD**: Standard Deviation; **AIC**: Akaike Information Criterion; **BIC**: Bayesian Information Criterion; **LogLik**: Log-likelihood.

### 5 Supplementary Table 5. Associations between serum marker concentrations and child neurometabolite ratios

#### 5.1 Maternal serum markers during pregnancy

##### 5.1.1 Child neurometabolite ratios in the midline parietal grey matter voxel

###### 5.1.1.1 Child glutamate ratios

| | | | | Linear regression with robust standard errors | | | | | | | | |
|---|---|---|---|---|---|---|---|---|---|---|---|---|
| **Participants** | | **Serum markers** | | **Unadjusted analysis** | | | | | **Adjusted analysis*** | | | |
| **n CHU** | **n CHEU** | **Marker type** | **Marker name** | **β** | **95% CI** | **SE** | **P-value** | **BH** | **β** | **95% CI** | **SE** | **P-value** |
| **40** | **34** | **Pro-inflammatory** | **GM-CSF** | -0.23 | -0.76 to 0.31 | 0.27 | 0.40 | 0.49 | | | | |
| | | | **IFN-γ** | -0.44 | -1.03 to 0.16 | 0.30 | 0.15 | 0.35 | | | | |
| | | | **IL-1β** | -0.52 | -1.14 to 0.09 | 0.31 | 0.09 | 0.26 | | | | |
| | | | **IL-2** | -0.35 | -0.82 to 0.12 | 0.23 | 0.14 | 0.47 | | | | |
| | | | **IL-5** | -0.72 | -1.61 to 0.17 | 0.45 | 0.11 | 0.14 | | | | |
| | | | **IL-6** | -0.31 | -0.74 to 0.12 | 0.21 | 0.15 | 0.19 | | | | |
| | | | **IL-7** | -0.78 | -1.74 to 0.18 | 0.48 | 0.11 | 0.12 | | | | |
| | | | **IL-8** | -0.24 | -0.88 to 0.39 | 0.32 | 0.45 | 0.79 | | | | |
| | | | **TNFα** | -0.32 | -1.30 to 0.66 | 0.49 | 0.52 | 0.76 | | | | |
| | | **Anti-inflammatory** | **IL-4** | -0.28 | -0.66 to 0.10 | 0.19 | 0.15 | 0.16 | | | | |
| | | | **IL-10** | -0.59 | -1.13 to -0.05 | 0.27 | 0.032 | 0.05 | | | | |
| | | | **IL-12p70** | -0.68 | -1.35 to 0.00 | 0.34 | 0.05 | 0.07 | | | | |
| | | | **IL-13** | **-0.44** | **-0.80 to -0.07** | **0.18** | **0.019** | **0.030** | **-0.41** | **-0.80 to -0.02** | **0.19** | **0.038** |
| | | **Monocyte activation** | **CD14** | -0.28 | -2.20 to 1.64 | 0.96 | 0.77 | 0.75 | | | | |
| | | | **CD163** | -0.05 | -1.11 to 1.02 | 0.53 | 0.93 | 0.93 | | | | |
| | | **Neuroinflammatory** | **NGAL** | 0.11 | -0.81 to 1.02 | 0.46 | 0.82 | 0.89 | | | | |
| | | | **MMP-9** | **-0.85** | **-1.58 to -0.13** | **0.36** | **0.022** | **0.044** | **-0.85** | **-1.57 to -0.12** | **0.36** | **0.023** |
| | | | **YKL-40** | -0.58 | -1.23 to 0.07 | 0.32 | 0.08 | 0.17 | | | | |

**CHU**: HIV-unexposed children; **CHEU**: HIV-exposed uninfected children; **β**: Effect size; **BH**: Benjamini-Hochberg corrected p-value. *Child age, child sex, and tissue composition.

###### 5.1.1.2 Child myo-inositol ratios

| | | | | Linear regression with robust standard errors | | | | | | | | |
|---|---|---|---|---|---|---|---|---|---|---|---|---|
| **Participants** | | **Serum markers** | | **Unadjusted analysis** | | | | | **Adjusted analysis*** | | | |
| **n CHU** | **n CHEU** | **Marker type** | **Marker name** | **β** | **95% CI** | **SE** | **P-value** | **BH** | **β** | **95% CI** | **SE** | **P-value** |
| **40** | **34** | **Pro-inflammatory** | **GM-CSF** | 0.03 | -0.45 to 0.51 | 0.24 | 0.91 | 0.99 | | | | |
| | | | **IFN-γ** | 0.04 | -0.59 to 0.67 | 0.32 | 0.90 | 0.97 | | | | |
| | | | **IL-1β** | 0.38 | -0.08 to 0.84 | 0.23 | 0.10 | 0.63 | | | | |
| | | | **IL-2** | 0.28 | -0.11 to 0.68 | 0.20 | 0.15 | 0.72 | | | | |
| | | | **IL-5** | **0.79** | **0.25 to 1.33** | **0.27** | **0.005** | **0.047** | **0.79** | **0.24 to 1.34** | **0.27** | **0.005** |
| | | | **IL-6** | 0.22 | -0.18 to 0.62 | 0.20 | 0.27 | 0.45 | | | | |
| | | | **IL-7** | 0.36 | -0.49 to 1.22 | 0.43 | 0.40 | 0.67 | | | | |
| | | | **IL-8** | 0.27 | -0.13 to 0.66 | 0.20 | 0.18 | 0.65 | | | | |
| | | | **TNFα** | 0.51 | -0.33 to 1.34 | 0.42 | 0.23 | 0.47 | | | | |
| | | **Anti-inflammatory** | **IL-4** | 0.19 | -0.14 to 0.53 | 0.17 | 0.25 | 0.54 | | | | |
| | | | **IL-10** | 0.28 | -0.29 to 0.86 | 0.29 | 0.33 | 0.41 | | | | |
| | | | **IL-12p70** | 0.38 | -0.19 to 0.95 | 0.29 | 0.19 | 0.57 | | | | |
| | | | **IL-13** | 0.11 | -0.29 to 0.50 | 0.20 | 0.59 | 0.91 | | | | |
| | | **Monocyte activation** | **CD14** | -0.37 | -1.61 to 0.86 | 0.62 | 0.55 | 0.59 | | | | |
| | | | **CD163** | -0.16 | -1.24 to 0.92 | 0.54 | 0.77 | 0.75 | | | | |
| | | **Neuroinflammatory** | **NGAL** | 0.16 | -0.89 to 1.21 | 0.53 | 0.77 | 0.77 | | | | |
| | | | **MMP-9** | -0.03 | -0.76 to 0.69 | 0.36 | 0.93 | 0.98 | | | | |
| | | | **YKL-40** | 0.18 | -0.71 to 1.07 | 0.45 | 0.69 | 0.66 | | | | |

**CHU**: HIV-unexposed children; **CHEU**: HIV-exposed uninfected children; **β**: Effect size; **BH**: Benjamini-Hochberg corrected p-value. *Child age, child sex, and tissue composition.

##### 5.1.1.3 Child N-acetyl-aspartate ratios

| | | | | Linear regression with robust standard errors | | | | | | | | |
|---|---|---|---|---|---|---|---|---|---|---|---|---|
| **Participants** | | **Serum markers** | | **Unadjusted analysis** | | | | | **Adjusted analysis*** | | | |
| **n CHU** | **n CHEU** | **Marker type** | **Marker name** | **β** | **95% CI** | **SE** | **P-value** | **BH** | **β** | **95% CI** | **SE** | **P-value** |
| **40** | **34** | **Pro-inflammatory** | **GM-CSF** | -0.16 | -0.67 to 0.34 | 0.25 | 0.52 | 0.65 | | | | |
| | | | **IFN-γ** | -0.15 | -0.70 to 0.40 | 0.28 | 0.59 | 0.81 | | | | |
| | | | **IL-1β** | -0.41 | -0.93 to 0.12 | 0.26 | 0.13 | 0.29 | | | | |
| | | | **IL-2** | -0.22 | -0.62 to 0.17 | 0.20 | 0.26 | 0.60 | | | | |
| | | | **IL-5** | -0.72 | -1.39 to -0.05 | 0.34 | **0.035** | 0.11 | | | | |
| | | | **IL-6** | -0.24 | -0.58 to 0.11 | 0.17 | 0.18 | 0.35 | | | | |
| | | | **IL-7** | -0.60 | -1.34 to 0.15 | 0.37 | 0.11 | 0.17 | | | | |
| | | | **IL-8** | -0.09 | -0.76 to 0.57 | 0.33 | 0.78 | 0.98 | | | | |
| | | | **TNFα** | -0.15 | -0.84 to 0.54 | 0.35 | 0.66 | 0.96 | | | | |
| | | **Anti-inflammatory** | **IL-4** | -0.23 | -0.60 to 0.15 | 0.19 | 0.23 | 0.24 | | | | |
| | | | **IL-10** | -0.47 | -0.90 to -0.03 | 0.22 | **0.036** | 0.10 | | | | |
| | | | **IL-12p70** | -0.66 | -1.16 to -0.16 | 0.25 | **0.011** | 0.07 | | | | |
| | | | **IL-13** | -0.25 | -0.56 to 0.06 | 0.15 | 0.11 | 0.22 | | | | |
| | | **Monocyte activation** | **CD14** | -0.25 | -2.28 to 1.79 | 1.02 | 0.81 | 0.77 | | | | |
| | | | **CD163** | -0.42 | -1.44 to 0.60 | 0.51 | 0.42 | 0.47 | | | | |
| | | **Neuroinflammatory** | **NGAL** | -0.32 | -1.18 to 0.54 | 0.43 | 0.46 | 0.72 | | | | |
| | | | **MMP-9** | **-1.01** | **-1.74 to -0.28** | **0.37** | **0.008** | **0.013** | **-1.01** | **-1.74 to -0.27** | **0.37** | **0.008** |
| | | | **YKL-40** | -0.40 | -1.08 to 0.27 | 0.34 | 0.23 | 0.55 | | | | |

**CHU**: HIV-unexposed children; **CHEU**: HIV-exposed uninfected children; **β**: Effect size; **BH**: Benjamini-Hochberg corrected p-value. *Child age, child sex, and tissue composition.

##### 5.1.2 Child neurometabolite ratios in the left parietal white matter voxel

###### 5.1.2.1 Child glutamate ratios

| | | | | Linear regression with robust standard errors | | | | | | | | |
|---|---|---|---|---|---|---|---|---|---|---|---|---|
| Participants | | Serum markers | | Unadjusted analysis | | | | | Adjusted analysis* | | | |
| n CHU | n CHEU | Marker type | Marker name | β | 95% CI | SE | P-value | BH | β | 95% CI | SE | P-value |
| 40 | 34 | Pro-inflammatory | GM-CSF | 0.06 | -0.55 to 0.67 | 0.31 | 0.84 | 0.90 | | | | |
| | | | IFN-γ | 0.05 | -0.57 to 0.67 | 0.31 | 0.86 | 0.86 | | | | |
| | | | IL-1β | -0.15 | -0.77 to 0.48 | 0.31 | 0.64 | 0.86 | | | | |
| | | | IL-2 | -0.05 | -0.54 to 0.44 | 0.25 | 0.84 | 0.91 | | | | |
| | | | IL-5 | -0.33 | -1.19 to 0.54 | 0.43 | 0.45 | 0.54 | | | | |
| | | | IL-6 | 0.08 | -0.34 to 0.50 | 0.21 | 0.72 | 0.82 | | | | |
| | | | IL-7 | -0.09 | -1.07 to 0.89 | 0.49 | 0.86 | 0.96 | | | | |
| | | | IL-8 | 0.01 | -0.50 to 0.53 | 0.26 | 0.96 | 0.95 | | | | |
| | | | TNFα | -0.35 | -1.26 to 0.55 | 0.46 | 0.44 | 0.56 | | | | |
| | | Anti-inflammatory | IL-4 | -0.03 | -0.39 to 0.34 | 0.18 | 0.89 | 0.94 | | | | |
| | | | IL-10 | -0.08 | -0.64 to 0.48 | 0.28 | 0.78 | 0.95 | | | | |
| | | | IL-12p70 | -0.03 | -0.75 to 0.69 | 0.36 | 0.93 | 1.00 | | | | |
| | | | IL-13 | -0.11 | -0.56 to 0.35 | 0.23 | 0.64 | 0.75 | | | | |
| | | Monocyte activation | CD14 | -0.27 | -2.55 to 2.00 | 1.14 | 0.81 | 0.82 | | | | |
| | | | CD163 | 0.32 | -0.85 to 1.49 | 0.59 | 0.59 | 0.85 | | | | |
| | | Neuroinflammatory | NGAL | 0.01 | -1.01 to 1.04 | 0.52 | 0.98 | 0.98 | | | | |
| | | | MMP-9 | -0.09 | -0.83 to 0.65 | 0.37 | 0.81 | 0.88 | | | | |
| | | | YKL-40 | -0.15 | -0.74 to 0.43 | 0.29 | 0.60 | 0.74 | | | | |

**CHU**: HIV-unexposed children; **CHEU**: HIV-exposed uninfected children; **β**: Effect size; **BH**: Benjamini-Hochberg corrected p-value. *Child age, child sex, and tissue composition.

##### 5.1.2.2 Child myo-inositol ratios

| Participants | | Serum markers | | Linear regression with robust standard errors | | | | | | | | |
|---|---|---|---|---|---|---|---|---|---|---|---|---|
| | | | | **Unadjusted analysis** | | | | | **Adjusted analysis*** | | | |
| **n CHU** | **n CHEU** | **Marker type** | **Marker name** | **β** | **95% CI** | **SE** | **P-value** | **BH** | **β** | **95% CI** | **SE** | **P-value** |
| **40** | **34** | **Pro-inflammatory** | **GM-CSF** | -0.10 | -0.59 to 0.40 | 0.25 | 0.70 | 0.68 | | | | |
| | | | **IFN-γ** | -0.03 | -0.60 to 0.55 | 0.29 | 0.92 | 0.92 | | | | |
| | | | **IL-1β** | 0.00 | -0.53 to 0.53 | 0.27 | 0.99 | 0.99 | | | | |
| | | | **IL-2** | -0.04 | -0.46 to 0.38 | 0.21 | 0.85 | 0.85 | | | | |
| | | | **IL-5** | 0.49 | -0.06 to 1.05 | 0.28 | 0.08 | 0.40 | | | | |
| | | | **IL-6** | 0.18 | -0.33 to 0.69 | 0.25 | 0.48 | 0.40 | | | | |
| | | | **IL-7** | 0.00 | -0.92 to 0.92 | 0.46 | 1.00 | 1.00 | | | | |
| | | | **IL-8** | 0.22 | -0.27 to 0.71 | 0.25 | 0.37 | 0.90 | | | | |
| | | | **TNFα** | 0.31 | -0.52 to 1.14 | 0.42 | 0.46 | 0.95 | | | | |
| | | **Anti-inflammatory** | **IL-4** | 0.03 | -0.33 to 0.38 | 0.18 | 0.89 | 0.91 | | | | |
| | | | **IL-10** | 0.21 | -0.34 to 0.76 | 0.27 | 0.45 | 0.71 | | | | |
| | | | **IL-12p70** | 0.23 | -0.26 to 0.73 | 0.25 | 0.35 | 0.77 | | | | |
| | | | **IL-13** | -0.02 | -0.41 to 0.37 | 0.20 | 0.91 | 0.90 | | | | |
| | | **Monocyte activation** | **CD14** | 0.58 | -0.81 to 1.96 | 0.69 | 0.41 | 0.43 | | | | |
| | | | **CD163** | 0.48 | -0.80 to 1.76 | 0.64 | 0.46 | 0.41 | | | | |
| | | **Neuroinflammatory** | **NGAL** | 0.36 | -0.49 to 1.21 | 0.42 | 0.40 | 0.49 | | | | |
| | | | **MMP-9** | 0.37 | -0.40 to 1.14 | 0.38 | 0.34 | 0.55 | | | | |
| | | | **YKL-40** | 0.40 | -0.54 to 1.34 | 0.47 | 0.40 | 0.26 | | | | |

**CHU**: HIV-unexposed children; **CHEU**: HIV-exposed uninfected children; **β**: Effect size; **BH**: Benjamini-Hochberg corrected p-value. *Child age, child sex, and tissue composition.

##### 5.1.3 Child neurometabolite ratios in the right parietal white matter voxel

###### 5.1.3.1 Child glutamate ratios

| | | | | Linear regression with robust standard errors | | | | | | | | |
|---|---|---|---|---|---|---|---|---|---|---|---|---|
| **Participants** | | **Serum markers** | | **Unadjusted analysis** | | | | | **Adjusted analysis*** | | | |
| **n CHU** | **n CHEU** | **Marker type** | **Marker name** | **β** | **95% CI** | **SE** | **P-value** | **BH** | **β** | **95% CI** | **SE** | **P-value** |
| **40** | **34** | **Pro-inflammatory** | **GM-CSF** | -0.09 | -0.55 to 0.38 | 0.24 | 0.72 | 0.96 | | | | |
| | | | **IFN-γ** | 0.03 | -0.56 to 0.63 | 0.30 | 0.91 | 0.90 | | | | |
| | | | **IL-1β** | -0.15 | -0.68 to 0.38 | 0.27 | 0.57 | 0.77 | | | | |
| | | | **IL-2** | -0.07 | -0.49 to 0.35 | 0.21 | 0.74 | 0.88 | | | | |
| | | | **IL-5** | -0.17 | -0.92 to 0.58 | 0.37 | 0.65 | 0.80 | | | | |
| | | | **IL-6** | 0.07 | -0.36 to 0.50 | 0.22 | 0.75 | 0.70 | | | | |
| | | | **IL-7** | -0.35 | -1.20 to 0.49 | 0.42 | 0.40 | 0.56 | | | | |
| | | | **IL-8** | 0.07 | -0.46 to 0.59 | 0.26 | 0.80 | 0.77 | | | | |
| | | | **TNFα** | 0.18 | -0.68 to 1.05 | 0.43 | 0.68 | 0.68 | | | | |
| | | **Anti-inflammatory** | **IL-4** | -0.01 | -0.41 to 0.40 | 0.20 | 0.98 | 0.98 | | | | |
| | | | **IL-10** | -0.42 | -0.92 to 0.07 | 0.25 | 0.09 | 0.12 | | | | |
| | | | **IL-12p70** | -0.31 | -0.80 to 0.19 | 0.25 | 0.22 | 0.44 | | | | |
| | | | **IL-13** | -0.03 | -0.49 to 0.43 | 0.23 | 0.90 | 0.88 | | | | |
| | | **Monocyte activation** | **CD14** | -0.29 | -1.88 to 1.30 | 0.80 | 0.72 | 0.67 | | | | |
| | | | **CD163** | 0.34 | -0.95 to 1.64 | 0.65 | 0.60 | 0.95 | | | | |
| | | **Neuroinflammatory** | **NGAL** | -0.44 | -1.33 to 0.45 | 0.45 | 0.33 | 0.64 | | | | |
| | | | **MMP-9** | -0.06 | -0.84 to 0.72 | 0.39 | 0.89 | 0.87 | | | | |
| | | | **YKL-40** | **-0.75** | **-1.37 to -0.14** | **0.31** | **0.018** | **0.032** | **-0.90** | **-1.47 to -0.33** | **0.29** | **0.002** |

**CHU**: HIV-unexposed children; **CHEU**: HIV-exposed uninfected children; **β**: Effect size; **BH**: Benjamini-Hochberg corrected p-value. *Child age, child sex, and tissue composition.

##### 5.1.3.2 Child myo-inositol ratios

| | | | | Linear regression with robust standard errors | | | | | | | | |
|---|---|---|---|---|---|---|---|---|---|---|---|---|
| **Participants** | | **Serum markers** | | **Unadjusted analysis** | | | | | **Adjusted analysis*** | | | |
| **n CHU** | **n CHEU** | **Marker type** | **Marker name** | **β** | **95% CI** | **SE** | **P-value** | **BH** | **β** | **95% CI** | **SE** | **P-value** |
| **40** | **34** | **Pro-inflammatory** | **GM-CSF** | -0.08 | -0.51 to 0.34 | 0.21 | 0.70 | 0.72 | | | | |
| | | | **IFN-γ** | -0.12 | -0.60 to 0.36 | 0.24 | 0.62 | 0.65 | | | | |
| | | | **IL-1β** | 0.03 | -0.42 to 0.48 | 0.23 | 0.91 | 0.92 | | | | |
| | | | **IL-2** | 0.00 | -0.42 to 0.42 | 0.21 | 0.99 | 0.99 | | | | |
| | | | **IL-5** | 0.38 | -0.17 to 0.93 | 0.28 | 0.18 | 0.41 | | | | |
| | | | **IL-6** | 0.38 | 0.09 to 0.68 | 0.15 | **0.012** | 0.09 | | | | |
| | | | **IL-7** | 0.03 | -0.83 to 0.88 | 0.43 | 0.95 | 0.94 | | | | |
| | | | **IL-8** | **0.62** | **0.11 to 1.14** | **0.26** | **0.018** | **0.009** | **0.64** | **0.10 to 1.17** | **0.27** | **0.020** |
| | | | **TNFα** | 0.73 | -0.16 to 1.62 | 0.45 | 0.11 | 0.25 | | | | |
| | | **Anti-inflammatory** | **IL-4** | 0.12 | -0.16 to 0.40 | 0.14 | 0.39 | 0.66 | | | | |
| | | | **IL-10** | 0.20 | -0.28 to 0.68 | 0.24 | 0.41 | 0.70 | | | | |
| | | | **IL-12p70** | 0.05 | -0.45 to 0.55 | 0.25 | 0.83 | 0.84 | | | | |
| | | | **IL-13** | 0.11 | -0.24 to 0.47 | 0.18 | 0.52 | 0.68 | | | | |
| | | **Monocyte activation** | **CD14** | -0.06 | -1.59 to 1.47 | 0.77 | 0.94 | 0.93 | | | | |
| | | | **CD163** | 0.44 | -0.53 to 1.40 | 0.48 | 0.37 | 0.59 | | | | |
| | | **Neuroinflammatory** | **NGAL** | 0.75 | 0.01 to 1.49 | 0.37 | **0.047** | 0.09 | | | | |
| | | | **MMP-9** | 0.54 | -0.12 to 1.19 | 0.33 | 0.11 | 0.17 | | | | |
| | | | **YKL-40** | 0.22 | -0.46 to 0.89 | 0.34 | 0.52 | 0.64 | | | | |

**CHU**: HIV-unexposed children; **CHEU**: HIV-exposed uninfected children; **β**: Effect size; **BH**: Benjamini-Hochberg corrected p-value. *Child age, child sex, and tissue composition.

##### 5.1.3.3 Child N-acetyl-aspartate ratios

| | | | | Linear regression with robust standard errors | | | | | | | | |
|---|---|---|---|---|---|---|---|---|---|---|---|---|
| **Participants** | | **Serum markers** | | **Unadjusted analysis** | | | | | **Adjusted analysis*** | | | |
| **n CHU** | **n CHEU** | **Marker type** | **Marker name** | **β** | **95% CI** | **SE** | **P-value** | **BH** | **β** | **95% CI** | **SE** | **P-value** |
| **40** | **34** | **Pro-inflammatory** | **GM-CSF** | -0.22 | -0.63 to 0.19 | 0.21 | 0.29 | 0.47 | | | | |
| | | | **IFN-γ** | 0.08 | -0.55 to 0.72 | 0.32 | 0.80 | 0.97 | | | | |
| | | | **IL-1β** | -0.15 | -0.70 to 0.41 | 0.28 | 0.60 | 0.58 | | | | |
| | | | **IL-2** | -0.04 | -0.49 to 0.41 | 0.23 | 0.84 | 0.83 | | | | |
| | | | **IL-5** | -0.38 | -1.01 to 0.24 | 0.31 | 0.23 | 0.74 | | | | |
| | | | **IL-6** | -0.33 | -0.71 to 0.06 | 0.19 | 0.10 | 0.10 | | | | |
| | | | **IL-7** | -0.13 | -0.91 to 0.65 | 0.39 | 0.74 | 0.83 | | | | |
| | | | **IL-8** | -0.21 | -0.92 to 0.50 | 0.35 | 0.56 | 0.77 | | | | |
| | | | **TNFα** | -0.30 | -0.99 to 0.40 | 0.35 | 0.40 | 0.94 | | | | |
| | | **Anti-inflammatory** | **IL-4** | -0.23 | -0.54 to 0.09 | 0.16 | 0.16 | 0.23 | | | | |
| | | | **IL-10** | -0.20 | -0.70 to 0.29 | 0.25 | 0.41 | 0.47 | | | | |
| | | | **IL-12p70** | -0.33 | -0.84 to 0.18 | 0.26 | 0.20 | 0.42 | | | | |
| | | | **IL-13** | -0.19 | -0.55 to 0.16 | 0.18 | 0.28 | 0.33 | | | | |
| | | **Monocyte activation** | **CD14** | -1.11 | -2.76 to 0.54 | 0.83 | 0.18 | 0.19 | | | | |
| | | | **CD163** | -0.52 | -1.92 to 0.87 | 0.70 | 0.46 | 0.37 | | | | |
| | | **Neuroinflammatory** | **NGAL** | -0.47 | -1.36 to 0.43 | 0.45 | 0.30 | 0.37 | | | | |
| | | | **MMP-9** | -0.70 | -1.55 to 0.15 | 0.43 | 0.11 | 0.13 | | | | |
| | | | **YKL-40** | -0.20 | -0.97 to 0.57 | 0.39 | 0.60 | 0.71 | | | | |

**CHU**: HIV-unexposed children; **CHEU**: HIV-exposed uninfected children; **β**: Effect size; **BH**: Benjamini-Hochberg corrected p-value. *Child age, child sex, and tissue composition.

#### 5.2 Infant serum markers at 6 weeks of age

##### 5.2.1 Child neurometabolite ratios in the midline parietal grey matter voxel

###### 5.2.1.1 Child glutamate ratios

| | | | | Linear regression with robust standard errors | | | | | | | | |
|---|---|---|---|---|---|---|---|---|---|---|---|---|
| Participants | | Serum markers | | Unadjusted analysis | | | | | Adjusted analysis* | | | |
| n CHU | n CHEU | Marker type | Marker name | β | 95% CI | SE | P-value | BH | β | 95% CI | SE | P-value |
| 29 | 23 | Pro-inflammatory | GM-CSF | -0.31 | -0.92 to 0.31 | 0.31 | 0.32 | 0.35 | | | | |
| | | | IFN-γ | -0.36 | -0.93 to 0.22 | 0.29 | 0.22 | 0.52 | | | | |
| | | | IL-1β | -0.31 | -1.00 to 0.38 | 0.34 | 0.38 | 0.89 | | | | |
| | | | IL-2 | -0.32 | -1.11 to 0.47 | 0.39 | 0.42 | 0.95 | | | | |
| | | | IL-5 | -0.25 | -1.40 to 0.90 | 0.57 | 0.66 | 0.99 | | | | |
| | | | IL-6 | -0.19 | -0.72 to 0.33 | 0.26 | 0.47 | 0.98 | | | | |
| | | | IL-7 | -0.64 | -1.57 to 0.30 | 0.47 | 0.18 | 0.25 | | | | |
| | | | IL-8 | -0.57 | -1.74 to 0.60 | 0.58 | 0.33 | 0.30 | | | | |
| | | | TNFα | 0.18 | -1.05 to 1.41 | 0.61 | 0.77 | 0.82 | | | | |
| | | Anti-inflammatory | IL-4 | -0.29 | -0.77 to 0.18 | 0.24 | 0.22 | 0.42 | | | | |
| | | | IL-10 | -0.22 | -0.89 to 0.46 | 0.34 | 0.52 | 0.92 | | | | |
| | | | IL-12p70 | -0.52 | -1.28 to 0.23 | 0.38 | 0.17 | 0.44 | | | | |
| | | | IL-13 | -0.25 | -0.66 to 0.17 | 0.21 | 0.24 | 0.85 | | | | |
| | | Monocyte activation | CD14 | 0.01 | -1.92 to 1.94 | 0.96 | 0.99 | 0.99 | | | | |
| | | | CD163 | 0.84 | -0.74 to 2.42 | 0.79 | 0.29 | 0.26 | | | | |
| | | Neuroinflammatory | NGAL | 1.28 | -0.38 to 2.93 | 0.82 | 0.13 | 0.13 | | | | |
| | | | MMP-9 | -0.37 | -0.99 to 0.24 | 0.31 | 0.23 | 0.60 | | | | |
| | | | YKL-40 | 0.20 | -1.20 to 1.61 | 0.70 | 0.77 | 0.80 | | | | |

**CHU**: HIV-unexposed children; **CHEU**: HIV-exposed uninfected children; **β**: Effect size; **BH**: Benjamini-Hochberg corrected p-value. *Child age, child sex, and tissue composition.

##### 5.2.1.2 Child myo-inositol ratios

| | | | | Linear regression with robust standard errors | | | | | | | | |
|---|---|---|---|---|---|---|---|---|---|---|---|---|
| **Participants** | | **Serum markers** | | **Unadjusted analysis** | | | | | **Adjusted analysis*** | | | |
| **n CHU** | **n CHEU** | **Marker type** | **Marker name** | **β** | **95% CI** | **SE** | **P-value** | **BH** | **β** | **95% CI** | **SE** | **P-value** |
| **29** | **23** | **Pro-inflammatory** | **GM-CSF** | -0.20 | -0.93 to 0.53 | 0.36 | 0.58 | 0.87 | | | | |
| | | | **IFN-γ** | 0.01 | -0.61 to 0.64 | 0.31 | 0.96 | 0.96 | | | | |
| | | | **IL-1β** | 0.30 | -0.17 to 0.76 | 0.23 | 0.21 | 0.48 | | | | |
| | | | **IL-2** | -0.12 | -0.71 to 0.47 | 0.29 | 0.68 | 0.92 | | | | |
| | | | **IL-5** | 0.03 | -0.74 to 0.81 | 0.38 | 0.93 | 0.93 | | | | |
| | | | **IL-6** | -0.02 | -0.45 to 0.41 | 0.21 | 0.94 | 0.98 | | | | |
| | | | **IL-7** | -0.43 | -1.21 to 0.35 | 0.39 | 0.28 | 0.54 | | | | |
| | | | **IL-8** | 0.29 | -0.29 to 0.87 | 0.29 | 0.31 | 0.83 | | | | |
| | | | **TNFα** | -0.05 | -1.11 to 1.02 | 0.53 | 0.93 | 0.94 | | | | |
| | | **Anti-inflammatory** | **IL-4** | 0.01 | -0.51 to 0.54 | 0.26 | 0.96 | 0.97 | | | | |
| | | | **IL-10** | -0.25 | -0.92 to 0.42 | 0.33 | 0.46 | 0.94 | | | | |
| | | | **IL-12p70** | -0.22 | -1.05 to 0.62 | 0.41 | 0.60 | 0.91 | | | | |
| | | | **IL-13** | -0.24 | -0.68 to 0.21 | 0.22 | 0.29 | 0.42 | | | | |
| | | **Monocyte activation** | **CD14** | -0.71 | -3.85 to 2.43 | 1.56 | 0.65 | 0.76 | | | | |
| | | | **CD163** | -0.85 | -1.81 to 0.11 | 0.48 | 0.08 | 0.25 | | | | |
| | | **Neuroinflammatory** | **NGAL** | -0.67 | -1.65 to 0.31 | 0.49 | 0.18 | 0.26 | | | | |
| | | | **MMP-9** | 0.38 | -0.07 to 0.83 | 0.22 | 0.10 | 0.34 | | | | |
| | | | **YKL-40** | -0.27 | -2.18 to 1.65 | 0.95 | 0.78 | 0.87 | | | | |

**CHU**: HIV-unexposed children; **CHEU**: HIV-exposed uninfected children; **β**: Effect size; **BH**: Benjamini-Hochberg corrected p-value. *Child age, child sex, and tissue composition.

##### 5.2.1.3 Child N-acetyl-aspartate ratios

| | | | | Linear regression with robust standard errors | | | | | | | | |
|---|---|---|---|---|---|---|---|---|---|---|---|---|
| **Participants** | | **Serum markers** | | **Unadjusted analysis** | | | | | **Adjusted analysis*** | | | |
| **n CHU** | **n CHEU** | **Marker type** | **Marker name** | **β** | **95% CI** | **SE** | **P-value** | **BH** | **β** | **95% CI** | **SE** | **P-value** |
| **29** | **23** | **Pro-inflammatory** | **GM-CSF** | 0.11 | -0.52 to 0.75 | 0.31 | 0.72 | 0.95 | | | | |
| | | | **IFN-γ** | 0.18 | -0.38 to 0.73 | 0.28 | 0.52 | 0.50 | | | | |
| | | | **IL-1β** | 0.05 | -0.56 to 0.65 | 0.30 | 0.88 | 0.90 | | | | |
| | | | **IL-2** | 0.15 | -0.52 to 0.82 | 0.33 | 0.65 | 0.86 | | | | |
| | | | **IL-5** | 0.31 | -0.60 to 1.22 | 0.45 | 0.49 | 0.77 | | | | |
| | | | **IL-6** | -0.12 | -0.58 to 0.34 | 0.23 | 0.59 | 0.84 | | | | |
| | | | **IL-7** | 0.21 | -0.74 to 1.16 | 0.47 | 0.65 | 0.58 | | | | |
| | | | **IL-8** | -0.48 | -1.62 to 0.66 | 0.57 | 0.40 | 0.43 | | | | |
| | | | **TNFα** | 0.00 | -1.24 to 1.23 | 0.61 | 1.00 | 1.00 | | | | |
| | | **Anti-inflammatory** | **IL-4** | -0.09 | -0.59 to 0.41 | 0.25 | 0.72 | 0.84 | | | | |
| | | | **IL-10** | 0.32 | -0.25 to 0.90 | 0.28 | 0.26 | 0.39 | | | | |
| | | | **IL-12p70** | 0.18 | -0.51 to 0.88 | 0.35 | 0.60 | 0.78 | | | | |
| | | | **IL-13** | -0.10 | -0.49 to 0.29 | 0.19 | 0.62 | 1.00 | | | | |
| | | **Monocyte activation** | **CD14** | 1.63 | 0.05 to 3.20 | 0.78 | **0.043** | 0.18 | | | | |
| | | | **CD163** | 1.10 | -0.28 to 2.48 | 0.69 | 0.12 | 0.08 | | | | |
| | | **Neuroinflammatory** | **NGAL** | 1.14 | 0.03 to 2.26 | 0.55 | **0.044** | 0.12 | | | | |
| | | | **MMP-9** | -0.08 | -0.77 to 0.60 | 0.34 | 0.81 | 0.99 | | | | |
| | | | **YKL-40** | 0.22 | -1.33 to 1.76 | 0.77 | 0.78 | 0.73 | | | | |

**CHU**: HIV-unexposed children; **CHEU**: HIV-exposed uninfected children; **β**: Effect size; **BH**: Benjamini-Hochberg corrected p-value. *Child age, child sex, and tissue composition.

##### 5.2.2 Child neurometabolite ratios in the left parietal white matter voxel

###### 5.2.2.1 Child glutamate ratios

| | | | | Linear regression with robust standard errors | | | | | | | | |
|---|---|---|---|---|---|---|---|---|---|---|---|---|
| **Participants** | | **Serum markers** | | **Unadjusted analysis** | | | | | **Adjusted analysis*** | | | |
| **n CHU** | **n CHEU** | **Marker type** | **Marker name** | **β** | **95% CI** | **SE** | **P-value** | **BH** | **β** | **95% CI** | **SE** | **P-value** |
| **29** | **23** | **Pro-inflammatory** | **GM-CSF** | -0.68 | -1.28 to -0.09 | 0.30 | **0.026** | 0.09 | | | | |
| | | | **IFN-γ** | -0.62 | -1.35 to 0.10 | 0.36 | 0.09 | 0.17 | | | | |
| | | | **IL-1β** | **-0.86** | **-1.37 to -0.35** | **0.25** | **0.001** | **0.015** | **-0.76** | **-1.34 to -0.18** | **0.29** | **0.011** |
| | | | **IL-2** | -0.60 | -1.55 to 0.35 | 0.47 | 0.21 | 0.25 | | | | |
| | | | **IL-5** | -0.81 | -1.63 to 0.01 | 0.41 | 0.05 | 0.13 | | | | |
| | | | **IL-6** | -0.36 | -1.03 to 0.31 | 0.33 | 0.29 | 0.41 | | | | |
| | | | **IL-7** | -0.86 | -1.92 to 0.21 | 0.53 | 0.11 | 0.20 | | | | |
| | | | **IL-8** | -0.61 | -1.57 to 0.34 | 0.47 | 0.20 | 0.43 | | | | |
| | | | **TNFα** | -0.06 | -1.22 to 1.11 | 0.58 | 0.92 | 0.98 | | | | |
| | | **Anti-inflammatory** | **IL-4** | -0.46 | -0.94 to 0.03 | 0.24 | 0.06 | 0.18 | | | | |
| | | | **IL-10** | -0.55 | -1.60 to 0.50 | 0.52 | 0.29 | 0.56 | | | | |
| | | | **IL-12p70** | -0.69 | -1.58 to 0.19 | 0.44 | 0.12 | 0.19 | | | | |
| | | | **IL-13** | -0.37 | -0.81 to 0.06 | 0.22 | 0.09 | 0.31 | | | | |
| | | **Monocyte activation** | **CD14** | 1.48 | -0.44 to 3.41 | 0.96 | 0.13 | 0.38 | | | | |
| | | | **CD163** | 0.66 | -0.95 to 2.27 | 0.80 | 0.41 | 0.62 | | | | |
| | | **Neuroinflammatory** | **NGAL** | -0.08 | -1.91 to 1.75 | 0.91 | 0.93 | 0.97 | | | | |
| | | | **MMP-9** | -0.62 | -1.43 to 0.19 | 0.40 | 0.13 | 0.25 | | | | |
| | | | **YKL-40** | -0.43 | -1.47 to 0.62 | 0.52 | 0.42 | 0.61 | | | | |

**CHU**: HIV-unexposed children; **CHEU**: HIV-exposed uninfected children; **β**: Effect size; **BH**: Benjamini-Hochberg corrected p-value. *Child age, child sex, and tissue composition.

##### 5.2.2.2 Child myo-inositol ratios

| | | | | Linear regression with robust standard errors | | | | | | | | |
|---|---|---|---|---|---|---|---|---|---|---|---|---|
| **Participants** | | **Serum markers** | | **Unadjusted analysis** | | | | | **Adjusted analysis*** | | | |
| **n CHU** | **n CHEU** | **Marker type** | **Marker name** | **β** | **95% CI** | **SE** | **P-value** | **BH** | **β** | **95% CI** | **SE** | **P-value** |
| **29** | **23** | **Pro-inflammatory** | **GM-CSF** | -0.19 | -0.76 to 0.38 | 0.28 | 0.50 | 0.69 | | | | |
| | | | **IFN-γ** | -0.17 | -0.75 to 0.41 | 0.29 | 0.56 | 0.59 | | | | |
| | | | **IL-1β** | -0.12 | -0.74 to 0.50 | 0.31 | 0.70 | 0.74 | | | | |
| | | | **IL-2** | -0.42 | -0.91 to 0.07 | 0.24 | 0.09 | 0.13 | | | | |
| | | | **IL-5** | -0.15 | -0.89 to 0.58 | 0.37 | 0.68 | 0.79 | | | | |
| | | | **IL-6** | -0.03 | -0.44 to 0.39 | 0.21 | 0.90 | 0.98 | | | | |
| | | | **IL-7** | -0.58 | -1.25 to 0.09 | 0.33 | 0.09 | 0.13 | | | | |
| | | | **IL-8** | -0.01 | -0.61 to 0.58 | 0.29 | 0.96 | 0.96 | | | | |
| | | | **TNFα** | -0.36 | -1.19 to 0.47 | 0.41 | 0.39 | 0.72 | | | | |
| | | **Anti-inflammatory** | **IL-4** | -0.04 | -0.40 to 0.33 | 0.18 | 0.84 | 0.83 | | | | |
| | | | **IL-10** | -0.30 | -1.24 to 0.63 | 0.46 | 0.51 | 0.70 | | | | |
| | | | **IL-12p70** | -0.32 | -0.89 to 0.24 | 0.28 | 0.25 | 0.26 | | | | |
| | | | **IL-13** | -0.18 | -0.62 to 0.26 | 0.22 | 0.41 | 0.32 | | | | |
| | | **Monocyte activation** | **CD14** | 0.41 | -1.49 to 2.30 | 0.94 | 0.67 | 0.86 | | | | |
| | | | **CD163** | -0.93 | -1.81 to -0.05 | 0.44 | **0.038** | 0.11 | | | | |
| | | **Neuroinflammatory** | **NGAL** | -0.88 | -1.76 to 0.00 | 0.44 | **0.049** | 0.19 | | | | |
| | | | **MMP-9** | -0.17 | -0.70 to 0.36 | 0.26 | 0.53 | 0.92 | | | | |
| | | | **YKL-40** | 0.11 | -2.25 to 2.47 | 1.17 | 0.93 | 0.99 | | | | |

**CHU**: HIV-unexposed children; **CHEU**: HIV-exposed uninfected children; **β**: Effect size; **BH**: Benjamini-Hochberg corrected p-value. *Child age, child sex, and tissue composition.

##### 5.2.3 Child neurometabolite ratios in the right parietal white matter voxel

###### 5.2.3.1 Child glutamate ratios

| | | | | Linear regression with robust standard errors | | | | | | | | |
|---|---|---|---|---|---|---|---|---|---|---|---|---|
| **Participants** | | **Serum markers** | | **Unadjusted analysis** | | | | | **Adjusted analysis*** | | | |
| **n CHU** | **n CHEU** | **Marker type** | **Marker name** | **β** | **95% CI** | **SE** | **P-value** | **BH** | **β** | **95% CI** | **SE** | **P-value** |
| **29** | **23** | **Pro-inflammatory** | **GM-CSF** | -0.38 | -0.89 to 0.13 | 0.25 | 0.14 | 0.29 | | | | |
| | | | **IFN-γ** | -0.51 | -1.16 to 0.13 | 0.32 | 0.12 | 0.13 | | | | |
| | | | **IL-1β** | -0.41 | -0.95 to 0.13 | 0.27 | 0.14 | 0.53 | | | | |
| | | | **IL-2** | -0.27 | -0.93 to 0.40 | 0.33 | 0.42 | 0.53 | | | | |
| | | | **IL-5** | -0.48 | -1.30 to 0.34 | 0.41 | 0.25 | 0.33 | | | | |
| | | | **IL-6** | -0.32 | -0.66 to 0.03 | 0.17 | 0.07 | 0.18 | | | | |
| | | | **IL-7** | -0.71 | -1.54 to 0.12 | 0.41 | 0.09 | 0.15 | | | | |
| | | | **IL-8** | 0.08 | -0.99 to 1.15 | 0.53 | 0.88 | 0.84 | | | | |
| | | | **TNFα** | 0.00 | -1.50 to 1.49 | 0.74 | 0.99 | 0.99 | | | | |
| | | **Anti-inflammatory** | **IL-4** | -0.34 | -0.81 to 0.12 | 0.23 | 0.14 | 0.19 | | | | |
| | | | **IL-10** | -0.19 | -1.21 to 0.83 | 0.51 | 0.71 | 0.68 | | | | |
| | | | **IL-12p70** | -0.49 | -1.24 to 0.25 | 0.37 | 0.19 | 0.23 | | | | |
| | | | **IL-13** | -0.22 | -0.65 to 0.20 | 0.21 | 0.29 | 0.48 | | | | |
| | | **Monocyte activation** | **CD14** | 1.62 | -0.60 to 3.83 | 1.10 | 0.15 | 0.24 | | | | |
| | | | **CD163** | 0.40 | -0.84 to 1.64 | 0.62 | 0.52 | 0.67 | | | | |
| | | **Neuroinflammatory** | **NGAL** | -0.36 | -1.68 to 0.96 | 0.66 | 0.59 | 0.73 | | | | |
| | | | **MMP-9** | -0.61 | -1.60 to 0.38 | 0.49 | 0.22 | 0.14 | | | | |
| | | | **YKL-40** | -0.27 | -1.73 to 1.20 | 0.73 | 0.71 | 0.67 | | | | |

**CHU**: HIV-unexposed children; **CHEU**: HIV-exposed uninfected children; **β**: Effect size; **BH**: Benjamini-Hochberg corrected p-value. *Child age, child sex, and tissue composition.

##### 5.2.3.2 Child myo-inositol ratios

| | | | | Linear regression with robust standard errors | | | | | | | | |
|---|---|---|---|---|---|---|---|---|---|---|---|---|
| **Participants** | | **Serum markers** | | **Unadjusted analysis** | | | | | **Adjusted analysis*** | | | |
| **n CHU** | **n CHEU** | **Marker type** | **Marker name** | **β** | **95% CI** | **SE** | **P-value** | **BH** | **β** | **95% CI** | **SE** | **P-value** |
| **29** | **23** | **Pro-inflammatory** | **GM-CSF** | -0.19 | -0.78 to 0.39 | -0.78 | 0.39 | 0.29 | | | | |
| | | | **IFN-γ** | -0.23 | -0.92 to 0.46 | -0.92 | 0.46 | 0.34 | | | | |
| | | | **IL-1β** | -0.04 | -0.49 to 0.41 | -0.49 | 0.41 | 0.22 | | | | |
| | | | **IL-2** | -0.21 | -0.79 to 0.37 | -0.79 | 0.37 | 0.29 | | | | |
| | | | **IL-5** | -0.10 | -0.69 to 0.49 | -0.69 | 0.49 | 0.29 | | | | |
| | | | **IL-6** | -0.20 | -0.56 to 0.16 | -0.56 | 0.16 | 0.18 | | | | |
| | | | **IL-7** | -0.43 | -1.33 to 0.46 | -1.33 | 0.46 | 0.45 | | | | |
| | | | **IL-8** | -0.02 | -0.88 to 0.83 | -0.88 | 0.83 | 0.42 | | | | |
| | | | **TNFα** | -0.18 | -1.28 to 0.93 | -1.28 | 0.93 | 0.55 | | | | |
| | | **Anti-inflammatory** | **IL-4** | -0.18 | -0.47 to 0.10 | -0.47 | 0.10 | 0.14 | | | | |
| | | | **IL-10** | -0.33 | -1.15 to 0.49 | -1.15 | 0.49 | 0.41 | | | | |
| | | | **IL-12p70** | -0.26 | -0.93 to 0.40 | -0.93 | 0.40 | 0.33 | | | | |
| | | | **IL-13** | -0.27 | -0.67 to 0.13 | -0.67 | 0.13 | 0.20 | | | | |
| | | **Monocyte activation** | **CD14** | 0.64 | -1.62 to 2.91 | -1.62 | 2.91 | 1.13 | | | | |
| | | | **CD163** | -0.56 | -1.77 to 0.64 | -1.77 | 0.64 | 0.60 | | | | |
| | | **Neuroinflammatory** | **NGAL** | -0.49 | -1.51 to 0.54 | -1.51 | 0.54 | 0.51 | | | | |
| | | | **MMP-9** | -0.04 | -0.79 to 0.71 | -0.79 | 0.71 | 0.37 | | | | |
| | | | **YKL-40** | -0.13 | -2.41 to 2.14 | -2.41 | 2.14 | 1.13 | | | | |

**CHU**: HIV-unexposed children; **CHEU**: HIV-exposed uninfected children; **β**: Effect size; **BH**: Benjamini-Hochberg corrected p-value. *Child age, child sex, and tissue composition.

##### 5.2.3.3 Child N-acetyl-aspartate ratios

| | | | | Linear regression with robust standard errors | | | | | | | | |
|---|---|---|---|---|---|---|---|---|---|---|---|---|
| **Participants** | | **Serum markers** | | **Unadjusted analysis** | | | | | **Adjusted analysis*** | | | |
| **n CHU** | **n CHEU** | **Marker type** | **Marker name** | **β** | **95% CI** | **SE** | **P-value** | **BH** | **β** | **95% CI** | **SE** | **P-value** |
| **29** | **23** | **Pro-inflammatory** | **GM-CSF** | -0.24 | -0.76 to 0.27 | 0.26 | 0.35 | 0.80 | | | | |
| | | | **IFN-γ** | -0.07 | -0.62 to 0.49 | 0.28 | 0.80 | 0.78 | | | | |
| | | | **IL-1β** | -0.15 | -0.72 to 0.42 | 0.28 | 0.59 | 0.53 | | | | |
| | | | **IL-2** | -0.13 | -0.76 to 0.51 | 0.32 | 0.69 | 0.63 | | | | |
| | | | **IL-5** | -0.14 | -0.89 to 0.60 | 0.37 | 0.70 | 0.85 | | | | |
| | | | **IL-6** | -0.22 | -0.62 to 0.18 | 0.20 | 0.27 | 0.28 | | | | |
| | | | **IL-7** | 0.02 | -0.89 to 0.94 | 0.45 | 0.96 | 0.95 | | | | |
| | | | **IL-8** | -0.04 | -1.30 to 1.21 | 0.62 | 0.94 | 0.90 | | | | |
| | | | **TNFα** | 0.82 | -0.25 to 1.89 | 0.53 | 0.13 | 0.08 | | | | |
| | | **Anti-inflammatory** | **IL-4** | -0.20 | -0.56 to 0.17 | 0.18 | 0.29 | 0.56 | | | | |
| | | | **IL-10** | 0.04 | -0.96 to 1.05 | 0.50 | 0.93 | 0.90 | | | | |
| | | | **IL-12p70** | -0.16 | -0.77 to 0.45 | 0.30 | 0.60 | 0.75 | | | | |
| | | | **IL-13** | -0.20 | -0.52 to 0.11 | 0.16 | 0.21 | 0.45 | | | | |
| | | **Monocyte activation** | **CD14** | -0.02 | -1.98 to 1.94 | 0.98 | 0.98 | 0.98 | | | | |
| | | | **CD163** | 1.20 | -0.17 to 2.57 | 0.68 | 0.08 | **0.047** | | | | |
| | | **Neuroinflammatory** | **NGAL** | 0.41 | -0.79 to 1.62 | 0.60 | 0.49 | 0.56 | | | | |
| | | | **MMP-9** | -0.11 | -0.68 to 0.45 | 0.28 | 0.69 | 0.90 | | | | |
| | | | **YKL-40** | 0.33 | -1.09 to 1.75 | 0.71 | 0.64 | 0.57 | | | | |

**CHU**: HIV-unexposed children; **CHEU**: HIV-exposed uninfected children; **β**: Effect size; **BH**: Benjamini-Hochberg corrected p-value. *Child age, child sex, and tissue composition.

##### 5.3 Child serum markers at 2 years of age

##### 5.3.1 Child neurometabolite ratios in the midline parietal grey matter voxel

###### 5.3.1.1 Child glutamate ratios

| | | | | Linear regression with robust standard errors | | | | | | | | |
|---|---|---|---|---|---|---|---|---|---|---|---|---|
| Participants | | Serum markers | | Unadjusted analysis | | | | | Adjusted analysis* | | | |
| n CHU | n CHEU | Marker type | Marker name | β | 95% CI | SE | P-value | BH | β | 95% CI | SE | P-value |
| 35 | 28 | Pro-inflammatory | GM-CSF | -0.17 | -0.86 to 0.51 | 0.34 | 0.61 | 0.98 | | | | |
| | | | IFN-γ | -0.01 | -0.90 to 0.89 | 0.45 | 0.99 | 0.99 | | | | |
| | | | IL-1β | -0.23 | -0.98 to 0.53 | 0.38 | 0.55 | 0.99 | | | | |
| | | | IL-2 | -0.76 | -1.56 to 0.03 | 0.40 | 0.06 | 0.16 | | | | |
| | | | IL-5 | 0.63 | -0.40 to 1.65 | 0.51 | 0.23 | 0.43 | | | | |
| | | | IL-6 | 0.10 | -0.62 to 0.83 | 0.36 | 0.77 | 0.89 | | | | |
| | | | IL-7 | -0.21 | -1.36 to 0.94 | 0.57 | 0.72 | 0.99 | | | | |
| | | | IL-8 | 0.25 | -0.28 to 0.79 | 0.27 | 0.35 | 0.82 | | | | |
| | | | TNFα | -0.43 | -1.39 to 0.53 | 0.48 | 0.37 | 0.56 | | | | |
| | | Anti-inflammatory | IL-4 | 0.01 | -0.56 to 0.57 | 0.28 | 0.98 | 1.00 | | | | |
| | | | IL-10 | -0.69 | -1.62 to 0.23 | 0.46 | 0.14 | 0.25 | | | | |
| | | | IL-12p70 | -0.68 | -1.49 to 0.14 | 0.41 | 0.10 | 0.24 | | | | |
| | | | IL-13 | 0.07 | -0.56 to 0.70 | 0.32 | 0.82 | 0.93 | | | | |
| | | Monocyte activation | CD14 | 1.13 | -0.17 to 2.43 | 0.65 | 0.09 | 0.28 | | | | |
| | | | CD163 | 0.52 | -0.74 to 1.78 | 0.63 | 0.41 | 0.61 | | | | |
| | | Neuroinflammatory | **NGAL** | **1.00** | **0.12 to 1.88** | **0.44** | **0.027** | **0.039** | **1.00** | **0.07 to 1.94** | **0.47** | **0.036** |
| | | | MMP-9 | 0.74 | -0.10 to 1.58 | 0.42 | 0.08 | 0.12 | | | | |
| | | | YKL-40 | 0.39 | -0.51 to 1.29 | 0.45 | 0.38 | 0.58 | | | | |

**CHU**: HIV-unexposed children; **CHEU**: HIV-exposed uninfected children; **β**: Effect size; **BH**: Benjamini-Hochberg corrected p-value. *Child age, child sex, and tissue composition.

##### 5.3.1.2 Child myo-inositol ratios

| | | | | Linear regression with robust standard errors | | | | | | | | |
|---|---|---|---|---|---|---|---|---|---|---|---|---|
| **Participants** | | **Serum markers** | | **Unadjusted analysis** | | | | | **Adjusted analysis*** | | | |
| **n CHU** | **n CHEU** | **Marker type** | **Marker name** | **β** | **95% CI** | **SE** | **P-value** | **BH** | **β** | **95% CI** | **SE** | **P-value** |
| **35** | **28** | **Pro-inflammatory** | **GM-CSF** | 0.03 | -0.51 to 0.57 | 0.27 | 0.90 | 0.92 | | | | |
| | | | **IFN-γ** | -0.01 | -0.90 to 0.88 | 0.44 | 0.98 | 0.98 | | | | |
| | | | **IL-1β** | 0.17 | -0.49 to 0.83 | 0.33 | 0.61 | 0.98 | | | | |
| | | | **IL-2** | 0.32 | -0.39 to 1.03 | 0.35 | 0.37 | 0.91 | | | | |
| | | | **IL-5** | -0.11 | -1.43 to 1.20 | 0.66 | 0.86 | 0.82 | | | | |
| | | | **IL-6** | -0.05 | -0.50 to 0.39 | 0.22 | 0.81 | 0.88 | | | | |
| | | | **IL-7** | -0.06 | -1.24 to 1.13 | 0.59 | 0.92 | 0.97 | | | | |
| | | | **IL-8** | 0.31 | -0.18 to 0.80 | 0.25 | 0.21 | 0.70 | | | | |
| | | | **TNFα** | 0.03 | -0.69 to 0.76 | 0.36 | 0.93 | 0.99 | | | | |
| | | **Anti-inflammatory** | **IL-4** | 0.17 | -0.25 to 0.59 | 0.21 | 0.41 | 0.94 | | | | |
| | | | **IL-10** | 0.16 | -0.83 to 1.16 | 0.50 | 0.74 | 0.77 | | | | |
| | | | **IL-12p70** | 0.24 | -0.53 to 1.01 | 0.39 | 0.53 | 0.73 | | | | |
| | | | **IL-13** | -0.41 | -0.93 to 0.12 | 0.26 | 0.13 | 0.31 | | | | |
| | | **Monocyte activation** | **CD14** | -0.52 | -1.72 to 0.67 | 0.60 | 0.39 | 0.51 | | | | |
| | | | **CD163** | -0.81 | -1.86 to 0.24 | 0.52 | 0.13 | 0.30 | | | | |
| | | **Neuroinflammatory** | **NGAL** | -0.41 | -1.16 to 0.33 | 0.37 | 0.27 | 0.71 | | | | |
| | | | **MMP-9** | **1.31** | **0.24 to 2.38** | **0.54** | **0.017** | **0.012** | **1.29** | **0.12 to 2.45** | **0.58** | **0.031** |
| | | | **YKL-40** | 0.46 | -0.56 to 1.47 | 0.51 | 0.37 | 0.70 | | | | |

**CHU**: HIV-unexposed children; **CHEU**: HIV-exposed uninfected children; **β**: Effect size; **BH**: Benjamini-Hochberg corrected p-value. *Child age, child sex, and tissue composition.

##### 5.3.1.3 Child N-acetyl-aspartate ratios

| | | | | Linear regression with robust standard errors | | | | | | | | |
|---|---|---|---|---|---|---|---|---|---|---|---|---|
| **Participants** | | **Serum markers** | | **Unadjusted analysis** | | | | | **Adjusted analysis*** | | | |
| **n CHU** | **n CHEU** | **Marker type** | **Marker name** | **β** | **95% CI** | **SE** | **P-value** | **BH** | **β** | **95% CI** | **SE** | **P-value** |
| **35** | **28** | **Pro-inflammatory** | **GM-CSF** | -0.02 | -0.57 to 0.53 | 0.28 | 0.94 | 0.94 | | | | |
| | | | **IFN-γ** | 0.08 | -0.79 to 0.94 | 0.43 | 0.86 | 0.85 | | | | |
| | | | **IL-1β** | -0.26 | -0.86 to 0.34 | 0.30 | 0.39 | 0.53 | | | | |
| | | | **IL-2** | -0.63 | -1.28 to 0.02 | 0.33 | 0.06 | 0.24 | | | | |
| | | | **IL-5** | -0.23 | -0.93 to 0.47 | 0.35 | 0.51 | 0.80 | | | | |
| | | | **IL-6** | -0.16 | -0.64 to 0.31 | 0.24 | 0.50 | 0.85 | | | | |
| | | | **IL-7** | -0.15 | -1.20 to 0.89 | 0.52 | 0.77 | 0.96 | | | | |
| | | | **IL-8** | -0.18 | -0.64 to 0.29 | 0.23 | 0.44 | 0.67 | | | | |
| | | | **TNFα** | -0.25 | -1.14 to 0.63 | 0.44 | 0.57 | 0.66 | | | | |
| | | **Anti-inflammatory** | **IL-4** | -0.27 | -0.69 to 0.16 | 0.21 | 0.21 | 0.30 | | | | |
| | | | **IL-10** | -0.68 | -1.50 to 0.13 | 0.41 | 0.10 | 0.24 | | | | |
| | | | **IL-12p70** | -0.65 | -1.36 to 0.07 | 0.36 | 0.07 | 0.15 | | | | |
| | | | **IL-13** | -0.15 | -0.60 to 0.31 | 0.23 | 0.52 | 0.72 | | | | |
| | | **Monocyte activation** | **CD14** | 0.78 | -0.57 to 2.13 | 0.68 | 0.25 | 0.55 | | | | |
| | | | **CD163** | 0.36 | -0.68 to 1.41 | 0.52 | 0.49 | 0.97 | | | | |
| | | **Neuroinflammatory** | **NGAL** | 0.80 | -0.13 to 1.73 | 0.47 | 0.09 | 0.09 | | | | |
| | | | **MMP-9** | 0.59 | -0.29 to 1.47 | 0.44 | 0.19 | 0.24 | | | | |
| | | | **YKL-40** | -0.21 | -1.02 to 0.59 | 0.40 | 0.60 | 0.81 | | | | |

**CHU**: HIV-unexposed children; **CHEU**: HIV-exposed uninfected children; **β**: Effect size; **BH**: Benjamini-Hochberg corrected p-value. *Child age, child sex, and tissue composition.

##### 5.3.2 Child neurometabolite ratios in the left parietal white matter voxel

###### 5.3.2.1 Child glutamate ratios

| | | | | Linear regression with robust standard errors | | | | | | | | |
|---|---|---|---|---|---|---|---|---|---|---|---|---|
| **Participants** | | **Serum markers** | | **Unadjusted analysis** | | | | | **Adjusted analysis*** | | | |
| **n CHU** | **n CHEU** | **Marker type** | **Marker name** | **β** | **95% CI** | **SE** | **P-value** | **BH** | **β** | **95% CI** | **SE** | **P-value** |
| **35** | **28** | **Pro-inflammatory** | **GM-CSF** | -0.12 | -0.84 to 0.60 | 0.36 | 0.74 | 0.97 | | | | |
| | | | **IFN-γ** | 0.14 | -0.63 to 0.91 | 0.38 | 0.72 | 0.94 | | | | |
| | | | **IL-1β** | -0.15 | -0.79 to 0.49 | 0.32 | 0.64 | 0.83 | | | | |
| | | | **IL-2** | -0.41 | -1.23 to 0.40 | 0.41 | 0.32 | 0.79 | | | | |
| | | | **IL-5** | 0.99 | 0.08 to 1.90 | 0.45 | 0.033 | 0.06 | | | | |
| | | | **IL-6** | 0.09 | -0.63 to 0.81 | 0.36 | 0.81 | 0.87 | | | | |
| | | | **IL-7** | 0.14 | -0.96 to 1.24 | 0.55 | 0.80 | 0.98 | | | | |
| | | | **IL-8** | 0.31 | -0.21 to 0.83 | 0.26 | 0.24 | 0.55 | | | | |
| | | | **TNFα** | -0.10 | -0.77 to 0.57 | 0.33 | 0.76 | 0.87 | | | | |
| | | **Anti-inflammatory** | **IL-4** | 0.09 | -0.36 to 0.54 | 0.22 | 0.70 | 0.89 | | | | |
| | | | **IL-10** | 0.09 | -1.11 to 1.29 | 0.60 | 0.88 | 0.92 | | | | |
| | | | **IL-12p70** | -0.24 | -1.23 to 0.75 | 0.49 | 0.63 | 0.96 | | | | |
| | | | **IL-13** | 0.18 | -0.31 to 0.67 | 0.25 | 0.47 | 0.98 | | | | |
| | | **Monocyte activation** | **CD14** | 0.41 | -1.20 to 2.01 | 0.80 | 0.62 | 0.59 | | | | |
| | | | **CD163** | 0.30 | -0.88 to 1.48 | 0.59 | 0.61 | 0.70 | | | | |
| | | **Neuroinflammatory** | **NGAL** | 0.47 | -0.36 to 1.29 | 0.41 | 0.26 | 0.39 | | | | |
| | | | **MMP-9** | -0.27 | -1.26 to 0.72 | 0.49 | 0.59 | 0.68 | | | | |
| | | | **YKL-40** | 0.37 | -0.44 to 1.18 | 0.40 | 0.37 | 0.42 | | | | |

**CHU**: HIV-unexposed children; **CHEU**: HIV-exposed uninfected children; **β**: Effect size; **BH**: Benjamini-Hochberg corrected p-value. *Child age, child sex, and tissue composition.

##### 5.3.2.2 Child myo-inositol ratios

| | | | | Linear regression with robust standard errors | | | | | | | | |
|---|---|---|---|---|---|---|---|---|---|---|---|---|
| **Participants** | | **Serum markers** | | **Unadjusted analysis** | | | | | **Adjusted analysis*** | | | |
| **n CHU** | **n CHEU** | **Marker type** | **Marker name** | **β** | **95% CI** | **SE** | **P-value** | **BH** | **β** | **95% CI** | **SE** | **P-value** |
| **35** | **28** | **Pro-inflammatory** | **GM-CSF** | 0.09 | -0.47 to 0.65 | 0.28 | 0.75 | 0.95 | | | | |
| | | | **IFN-γ** | -0.36 | -1.32 to 0.60 | 0.48 | 0.46 | 0.35 | | | | |
| | | | **IL-1β** | -0.09 | -0.75 to 0.57 | 0.33 | 0.78 | 0.78 | | | | |
| | | | **IL-2** | 0.12 | -0.58 to 0.83 | 0.35 | 0.73 | 0.74 | | | | |
| | | | **IL-5** | -0.05 | -0.87 to 0.77 | 0.41 | 0.90 | 0.91 | | | | |
| | | | **IL-6** | -0.14 | -0.65 to 0.37 | 0.26 | 0.59 | 0.75 | | | | |
| | | | **IL-7** | -0.88 | -2.14 to 0.39 | 0.63 | 0.17 | 0.13 | | | | |
| | | | **IL-8** | 0.06 | -0.41 to 0.53 | 0.23 | 0.79 | 0.82 | | | | |
| | | | **TNFα** | -0.53 | -1.40 to 0.33 | 0.43 | 0.22 | 0.34 | | | | |
| | | **Anti-inflammatory** | **IL-4** | -0.07 | -0.55 to 0.41 | 0.24 | 0.76 | 0.75 | | | | |
| | | | **IL-10** | 0.11 | -0.73 to 0.95 | 0.42 | 0.79 | 0.96 | | | | |
| | | | **IL-12p70** | 0.01 | -0.71 to 0.74 | 0.36 | 0.97 | 0.97 | | | | |
| | | | **IL-13** | -0.42 | -0.98 to 0.15 | 0.28 | 0.15 | 0.17 | | | | |
| | | **Monocyte activation** | **CD14** | 0.10 | -1.26 to 1.46 | 0.68 | 0.88 | 0.94 | | | | |
| | | | **CD163** | -0.60 | -1.56 to 0.36 | 0.48 | 0.22 | 0.52 | | | | |
| | | **Neuroinflammatory** | **NGAL** | 0.17 | -0.67 to 1.01 | 0.42 | 0.69 | 0.79 | | | | |
| | | | **MMP-9** | 0.79 | -0.06 to 1.64 | 0.42 | 0.07 | 0.20 | | | | |
| | | | **YKL-40** | 0.72 | -0.21 to 1.64 | 0.46 | 0.13 | 0.21 | | | | |

**CHU**: HIV-unexposed children; **CHEU**: HIV-exposed uninfected children; **β**: Effect size; **BH**: Benjamini-Hochberg corrected p-value. *Child age, child sex, and tissue composition.

##### 5.3.3 Child neurometabolite ratios in the right parietal white matter voxel

###### 5.3.3.1 Child glutamate ratios

| | | | | Linear regression with robust standard errors | | | | | | | | |
|---|---|---|---|---|---|---|---|---|---|---|---|---|
| **Participants** | | **Serum markers** | | **Unadjusted analysis** | | | | | **Adjusted analysis*** | | | |
| **n CHU** | **n CHEU** | **Marker type** | **Marker name** | **β** | **95% CI** | **SE** | **P-value** | **BH** | **β** | **95% CI** | **SE** | **P-value** |
| **35** | **28** | **Pro-inflammatory** | **GM-CSF** | -0.39 | -0.91 to 0.14 | 0.26 | 0.14 | 0.26 | | | | |
| | | | **IFN-γ** | 0.08 | -0.76 to 0.93 | 0.42 | 0.84 | 0.82 | | | | |
| | | | **IL-1β** | -0.23 | -0.89 to 0.43 | 0.33 | 0.49 | 0.61 | | | | |
| | | | **IL-2** | -0.24 | -0.87 to 0.40 | 0.32 | 0.46 | 0.65 | | | | |
| | | | **IL-5** | 0.56 | -0.33 to 1.46 | 0.45 | 0.21 | 0.58 | | | | |
| | | | **IL-6** | 0.05 | -0.53 to 0.64 | 0.29 | 0.85 | 0.94 | | | | |
| | | | **IL-7** | -0.09 | -1.38 to 1.20 | 0.64 | 0.89 | 0.87 | | | | |
| | | | **IL-8** | 0.27 | -0.25 to 0.79 | 0.26 | 0.30 | 0.50 | | | | |
| | | | **TNFα** | -0.15 | -1.04 to 0.74 | 0.45 | 0.73 | 0.67 | | | | |
| | | **Anti-inflammatory** | **IL-4** | -0.03 | -0.59 to 0.53 | 0.28 | 0.93 | 0.91 | | | | |
| | | | **IL-10** | -0.44 | -1.23 to 0.35 | 0.39 | 0.27 | 0.56 | | | | |
| | | | **IL-12p70** | -0.36 | -1.01 to 0.28 | 0.32 | 0.26 | 0.61 | | | | |
| | | | **IL-13** | 0.16 | -0.42 to 0.74 | 0.29 | 0.59 | 0.76 | | | | |
| | | **Monocyte activation** | **CD14** | 0.46 | -1.15 to 2.08 | 0.81 | 0.57 | 0.58 | | | | |
| | | | **CD163** | 0.49 | -0.50 to 1.47 | 0.49 | 0.33 | 0.75 | | | | |
| | | **Neuroinflammatory** | **NGAL** | 0.63 | -0.20 to 1.46 | 0.42 | 0.13 | 0.18 | | | | |
| | | | **MMP-9** | 0.16 | -0.66 to 0.98 | 0.41 | 0.70 | 0.87 | | | | |
| | | | **YKL-40** | -0.11 | -0.87 to 0.65 | 0.38 | 0.77 | 0.79 | | | | |

**CHU**: HIV-unexposed children; **CHEU**: HIV-exposed uninfected children; **β**: Effect size; **BH**: Benjamini-Hochberg corrected p-value. *Child age, child sex, and tissue composition.

##### 5.3.3.2 Child myo-inositol ratios

| | | | | Linear regression with robust standard errors | | | | | | | | |
|---|---|---|---|---|---|---|---|---|---|---|---|---|
| Participants | | Serum markers | | Unadjusted analysis | | | | | Adjusted analysis* | | | |
| n CHU | n CHEU | Marker type | Marker name | β | 95% CI | SE | P-value | BH | β | 95% CI | SE | P-value |
| 35 | 28 | **Pro-inflammatory** | **GM-CSF** | 0.10 | -0.40 to 0.61 | 0.25 | 0.69 | 0.91 | | | | |
| | | | **IFN-γ** | -0.41 | -1.14 to 0.32 | 0.37 | 0.27 | 0.28 | | | | |
| | | | **IL-1β** | -0.14 | -0.72 to 0.45 | 0.29 | 0.64 | 0.67 | | | | |
| | | | **IL-2** | 0.01 | -0.61 to 0.62 | 0.31 | 0.98 | 0.98 | | | | |
| | | | **IL-5** | -0.25 | -1.21 to 0.72 | 0.48 | 0.61 | 0.78 | | | | |
| | | | **IL-6** | -0.01 | -0.51 to 0.49 | 0.25 | 0.96 | 0.95 | | | | |
| | | | **IL-7** | -0.72 | -1.78 to 0.34 | 0.53 | 0.18 | 0.20 | | | | |
| | | | **IL-8** | -0.15 | -0.71 to 0.40 | 0.28 | 0.59 | 0.58 | | | | |
| | | | **TNFα** | -0.62 | -1.34 to 0.10 | 0.36 | 0.09 | 0.14 | | | | |
| | | **Anti-inflammatory** | **IL-4** | -0.08 | -0.47 to 0.32 | 0.20 | 0.70 | 0.74 | | | | |
| | | | **IL-10** | -0.24 | -1.04 to 0.57 | 0.40 | 0.56 | 0.66 | | | | |
| | | | **IL-12p70** | -0.08 | -0.74 to 0.58 | 0.33 | 0.81 | 0.84 | | | | |
| | | | **IL-13** | -0.37 | -0.92 to 0.18 | 0.28 | 0.19 | 0.19 | | | | |
| | | **Monocyte activation** | **CD14** | -0.08 | -1.45 to 1.29 | 0.69 | 0.91 | 0.91 | | | | |
| | | | **CD163** | -0.53 | -1.64 to 0.59 | 0.56 | 0.35 | 0.62 | | | | |
| | | **Neuroinflammatory** | **NGAL** | 0.37 | -0.48 to 1.21 | 0.42 | 0.39 | 0.48 | | | | |
| | | | **MMP-9** | 0.72 | -0.32 to 1.75 | 0.52 | 0.17 | 0.33 | | | | |
| | | | **YKL-40** | 0.72 | -0.27 to 1.71 | 0.49 | 0.15 | 0.19 | | | | |

**CHU**: HIV-unexposed children; **CHEU**: HIV-exposed uninfected children; **β**: Effect size; **BH**: Benjamini-Hochberg corrected p-value. *Child age, child sex, and tissue composition.

###### 5.3.3.3 Child N-acetyl-aspartate ratios

| | | | | Linear regression with robust standard errors | | | | | | | | |
|---|---|---|---|---|---|---|---|---|---|---|---|---|
| **Participants** | | **Serum markers** | | **Unadjusted analysis** | | | | | **Adjusted analysis*** | | | |
| **n CHU** | **n CHEU** | **Marker type** | **Marker name** | **β** | **95% CI** | **SE** | **P-value** | **BH** | **β** | **95% CI** | **SE** | **P-value** |
| **35** | **28** | **Pro-inflammatory** | **GM-CSF** | -0.08 | -0.55 to 0.38 | 0.23 | 0.72 | 0.94 | | | | |
| | | | **IFN-γ** | 0.13 | -0.85 to 1.11 | 0.49 | 0.79 | 0.91 | | | | |
| | | | **IL-1β** | -0.12 | -0.82 to 0.57 | 0.35 | 0.72 | 0.73 | | | | |
| | | | **IL-2** | -0.28 | -1.05 to 0.49 | 0.39 | 0.47 | 0.49 | | | | |
| | | | **IL-5** | -0.05 | -0.90 to 0.81 | 0.43 | 0.91 | 0.97 | | | | |
| | | | **IL-6** | 0.06 | -0.49 to 0.61 | 0.28 | 0.82 | 0.79 | | | | |
| | | | **IL-7** | 0.28 | -1.11 to 1.67 | 0.70 | 0.69 | 0.88 | | | | |
| | | | **IL-8** | -0.11 | -0.60 to 0.38 | 0.24 | 0.65 | 0.86 | | | | |
| | | | **TNFα** | 0.45 | -0.47 to 1.38 | 0.46 | 0.33 | 0.38 | | | | |
| | | **Anti-inflammatory** | **IL-4** | 0.03 | -0.57 to 0.62 | 0.30 | 0.93 | 0.90 | | | | |
| | | | **IL-10** | -0.31 | -1.19 to 0.58 | 0.44 | 0.49 | 0.69 | | | | |
| | | | **IL-12p70** | -0.33 | -1.07 to 0.40 | 0.37 | 0.37 | 0.69 | | | | |
| | | | **IL-13** | 0.05 | -0.72 to 0.81 | 0.38 | 0.90 | 0.84 | | | | |
| | | **Monocyte activation** | **CD14** | -0.81 | -2.15 to 0.53 | 0.67 | 0.23 | 0.44 | | | | |
| | | | **CD163** | 0.13 | -0.86 to 1.13 | 0.50 | 0.79 | 0.79 | | | | |
| | | **Neuroinflammatory** | **NGAL** | 0.33 | -0.54 to 1.21 | 0.44 | 0.45 | 0.40 | | | | |
| | | | **MMP-9** | -0.24 | -1.09 to 0.61 | 0.43 | 0.57 | 0.70 | | | | |
| | | | **YKL-40** | -0.54 | -1.28 to 0.20 | 0.37 | 0.15 | 0.46 | | | | |

**CHU**: HIV-unexposed children; **CHEU**: HIV-exposed uninfected children; **β**: Effect size; **BH**: Benjamini-Hochberg corrected p-value. *Child age, child sex, and tissue composition.

#### 6 Supplementary Table 6. Sensitivity analyses

| | | | | Adjusted analysis[A] | | | Maternal age at delivery[B] | | | Maternal depression[C] | | | Maternal alcohol use[D] | | |
|---|---|---|---|---|---|---|---|---|---|---|---|---|---|---|---|
| **Timepoint** | **Brain region** | **Metabolite ratios** | **Serum marker** | **β** | **95% CI** | **P-value** | **β** | **95% CI** | **P-value** | **β** | **95% CI** | **P-value** | **β** | **95% CI** | **P-value** |
| **Pregnancy** | **PGM** | Glu | IL-13 | -0.41 | -0.80 to -0.02 | **0.038** | -0.39 | -0.75 to -0.05 | **0.031** | -0.46 | -0.83 to -0.09 | **0.016** | -0.50 | -0.89 to -0.10 | **0.016** |
| | | | MMP-9 | -0.85 | -1.57 to -0.12 | **0.023** | -0.81 | -1.57 to -0.04 | **0.039** | -0.87 | -1.63 to -0.11 | **0.026** | -0.88 | -1.63 to -0.13 | **0.023** |
| | | Myo | IL-5 | 0.79 | 0.24 to 1.34 | **0.005** | 0.78 | 0.25 to 1.32 | **0.005** | 0.80 | 0.26 to 1.35 | **0.005** | 0.78 | 0.26 to 1.30 | **0.004** |
| | | NAA | MMP-9 | -1.01 | -1.74 to -0.27 | **0.008** | -0.99 | -1.71 to -0.27 | **0.008** | -1.03 | -1.77 to -0.29 | **0.007** | -1.09 | -1.79 to -0.40 | **0.003** |
| | **RPWM** | Glu | YKL-40 | -0.90 | -1.47 to -0.33 | **0.002** | -0.90 | -1.46 to -0.35 | **0.002** | -0.91 | -1.47 to -0.34 | **0.002** | -0.91 | -1.52 to -0.31 | **0.004** |
| | | Myo | IL-8 | 0.64 | 0.10 to 1.17 | **0.020** | 0.64 | 0.10 to 1.19 | **0.021** | 0.64 | 0.08 to 1.20 | **0.025** | 0.64 | 0.10 to 1.18 | **0.021** |
| | **All regions** | Myo* | IL-5 | 0.84 | 0.23 to 1.44 | **0.007** | 0.81 | 0.20 to 1.42 | **0.010** | 0.86 | 0.24 to 1.47 | **0.007** | 0.83 | 0.25 to 1.40 | **0.006** |
| **6 weeks** | **LPWM** | Glu | IL-1β | -0.76 | -1.34 to -0.18 | **0.011** | -0.76 | -1.32 t -0.21 | **0.008** | -0.76 | -1.31 t -0.21 | **0.008** | -0.76 | -1.31 to -0.21 | **0.008** |
| **2 years** | **PGM** | Glu | NGAL | 1.00 | 0.07 to 1.94 | **0.036** | 1.06 | 0.20 to 1.93 | **0.017** | 1.20 | 0.27 to 2.14 | **0.013** | 1.05 | 0.12 to 1.98 | **0.027** |
| | | Myo | MMP-9 | 1.29 | 0.12 to 2.45 | **0.031** | 1.29 | 0.28 to 2.30 | **0.014** | 1.26 | 0.23 to 2.30 | **0.018** | 1.31 | 0.29 to 2.34 | **0.013** |
| | **L & RPWM** | Glu* | NGAL | 0.84 | 0.14 to 1.54 | **0.019** | 0.85 | 0.16 to 1.54 | **0.016** | 0.94 | 0.24 to 1.63 | **0.009** | 0.89 | 0.20 to 1.58 | **0.012** |

Sensitivity analyses of statistically significant associations between maternal and child immune marker concentrations and child neurometabolite ratios in children who are HIV-exposed and uninfected compared to HIV-unexposed peers. Linear regression models with robust standard errors were constructed including the following covariates:

A. Child age, child sex, and voxel tissue composition (reference)
B. Child age, child sex, and maternal age at delivery (sensitivity analysis B)
C. Child age, child sex, and antenatal maternal depression (sensitivity analysis C) (Note: missing data were handled using the last observation carried forward (LOCF) method)
D. Child age, child sex, and maternal alcohol use during pregnancy (sensitivity analysis D)

**MRS**: Magnetic Resonance Spectroscopy; **β**: Effect size; **PGM**: Parietal Grey Matter; **LPWM**: Left Parietal White Matter; **RPWM**: Right Parietal White Matter.

*Cross-regional neurometabolite patterns (factor loadings) previously reported in the same cohort, identified with factor analysis.

#### 7 Supplementary Table 7. Mediation analyses

Structural equation modelling estimates for direct, indirect, and total effects of maternal HIV on child neurometabolite ratios mediated by serum marker concentrations

| Timepoint | n CHU | n CHEU | Brain region | Neurometabolite ratios | Marker | Effect | Estimate | StdEst | 95% CI | SE | z-value | p-value |
|---|---|---|---|---|---|---|---|---|---|---|---|---|
| **Pregnancy** | **40** | **34** | **PGM** | Glutamate | IL-13 | Direct | 0.02 | 0.01 | -0.48 to 0.50 | 0.25 | 0.08 | 0.94 |
| | | | | | | Indirect | -0.02 | -0.01 | -0.15 to 0.12 | 0.07 | -0.30 | 0.76 |
| | | | | | | Total | ≈0.00 | ≈0.00 | -0.47 to 0.48 | 0.25 | ≈0.00 | 1.00 |
| | | | | | MMP-9 | Direct | -0.11 | -0.05 | -0.61 to 0.40 | 0.26 | -0.41 | 0.69 |
| | | | | | | Indirect | 0.11 | 0.05 | -0.07 to 0.28 | 0.09 | 1.16 | 0.25 |
| | | | | | | Total | ≈0.00 | ≈0.00 | | 0.26 | ≈0.00 | 1.00 |
| | | | | Myo-inositol | IL-5 | Direct | 0.09 | 0.05 | -0.38 to 0.51 | 0.22 | 0.40 | 0.69 |
| | | | | | | Indirect | 0.01 | 0.01 | -0.07 to 0.13 | 0.05 | 0.26 | 0.80 |
| | | | | | | Total | 0.10 | 0.05 | -0.35 to 0.53 | 0.23 | 0.44 | 0.66 |
| | | | | N-acetyl-aspartate | MMP-9 | Direct | -0.28 | -0.14 | -0.75 to 0.28 | 0.27 | -1.06 | 0.29 |
| | | | | | | Indirect | 0.05 | 0.02 | -0.12 to 0.23 | 0.09 | 0.56 | 0.58 |
| | | | | | | Total | -0.24 | -0.12 | -0.68 to 0.27 | 0.24 | -0.99 | 0.33 |
| | | | **RPWM** | Glutamate | YKL-40 | Direct | 0.59 | 0.29 | 0.11 to 1.05 | 0.24 | 2.48 | **0.013** |
| | | | | | | Indirect | 0.01 | ≈0.00 | -0.07 to 0.09 | 0.04 | 0.23 | 0.82 |
| | | | | | | Total | 0.60 | 0.29 | 0.15 to 1.06 | 0.23 | 2.54 | **0.011** |
| | | | | Myo-inositol | IL-8 | Direct | 0.62 | 0.32 | 0.22 to 1.04 | 0.21 | 2.94 | **0.003** |
| | | | | | | Indirect | 0.00 | 0.00 | -0.05 to 0.10 | 0.03 | 0.06 | 0.95 |
| | | | | | | Total | 0.63 | 0.32 | 0.24 to 1.05 | 0.21 | 2.99 | **0.003** |
| | | | **All regions** | Myo-inositol pattern | IL-5 | Direct | 0.55 | 0.24 | 0.04 to 1.05 | 0.26 | 2.12 | **0.034** |
| | | | | | | Indirect | ≈0.00 | ≈0.00 | -0.14 to 0.13 | 0.07 | -0.06 | 0.95 |
| | | | | | | Total | 0.54 | 0.23 | 0.04 to 1.05 | 0.26 | 2.10 | **0.036** |

| | | | | | | | | | | | | |
|---|---|---|---|---|---|---|---|---|---|---|---|---|
| **6 weeks** | **29** | **23** | **LPWM** | Glutamate | IL-1β | Direct | 0.06 | 0.03 | -0.54 to 0.65 | 0.30 | 0.21 | 0.83 |
| | | | | | | Indirect | ≈0.00 | ≈0.00 | -0.34 to 0.23 | 0.05 | 0.07 | 0.95 |
| | | | | | | Total | 0.07 | 0.03 | -0.54 to 0.65 | 0.30 | 0.23 | 0.82 |
| **2 years** | **35** | **28** | **PGM** | Glutamate | NGAL | Direct | -0.04 | -0.02 | -0.59 to 0.46 | 0.27 | -0.14 | 0.89 |
| | | | | | | Indirect | ≈0.00 | ≈0.00 | -0.07 to 0.08 | 0.04 | 0.08 | 0.94 |
| | | | | | | Total | -0.04 | -0.02 | -0.57 to 0.48 | 0.27 | -0.13 | 0.90 |
| | | | | Myo-inositol | MMP-9 | Direct | 0.03 | 0.02 | -0.45 to 0.51 | 0.25 | 0.12 | 0.90 |
| | | | | | | Indirect | 0.01 | ≈0.00 | -0.06 to 0.08 | 0.02 | 0.49 | 0.62 |
| | | | | | | Total | 0.04 | 0.02 | -0.42 to 0.51 | 0.24 | 0.14 | 0.89 |
| | | | **L & RPWM** | Glutamate pattern | NGAL | Direct | 0.24 | 0.14 | -0.17 to 0.65 | 0.21 | 1.18 | 0.24 |
| | | | | | | Indirect | ≈0.00 | ≈0.00 | -0.09 to 0.10 | 0.05 | 0.15 | 0.88 |
| | | | | | | Total | 0.25 | 0.15 | -0.16 to 0.64 | 0.20 | 1.25 | 0.21 |

**Predictor**: maternal HIV status; **Mediator**: maternal/child serum marker concentrations; **Outcome**: child neurometabolite ratios to total creatine at age 2–3 years.

**Direct path**: Predictor → Outcome
**Indirect path**: Predictor → Mediator → Outcome
**Total path**: Predictor → (Mediator + Outcome)

**CHEU**: Children who are HIV-Exposed and Uninfected; **CHU**: Children who are HIV-Unexposed; **MRS**: Magnetic Resonance Spectroscopy; **PGM**: Parietal Grey Matter; **LPWM**: Left Parietal White Matter; **RPWM**: Right Parietal White Matter; **Estimate**: Raw unstandardized regression coefficient; **StdEst**: Standardized regression coefficient for interpretability; **SE**: Standard Error.

#### 8 Supplementary Figure 1. Sample sizes

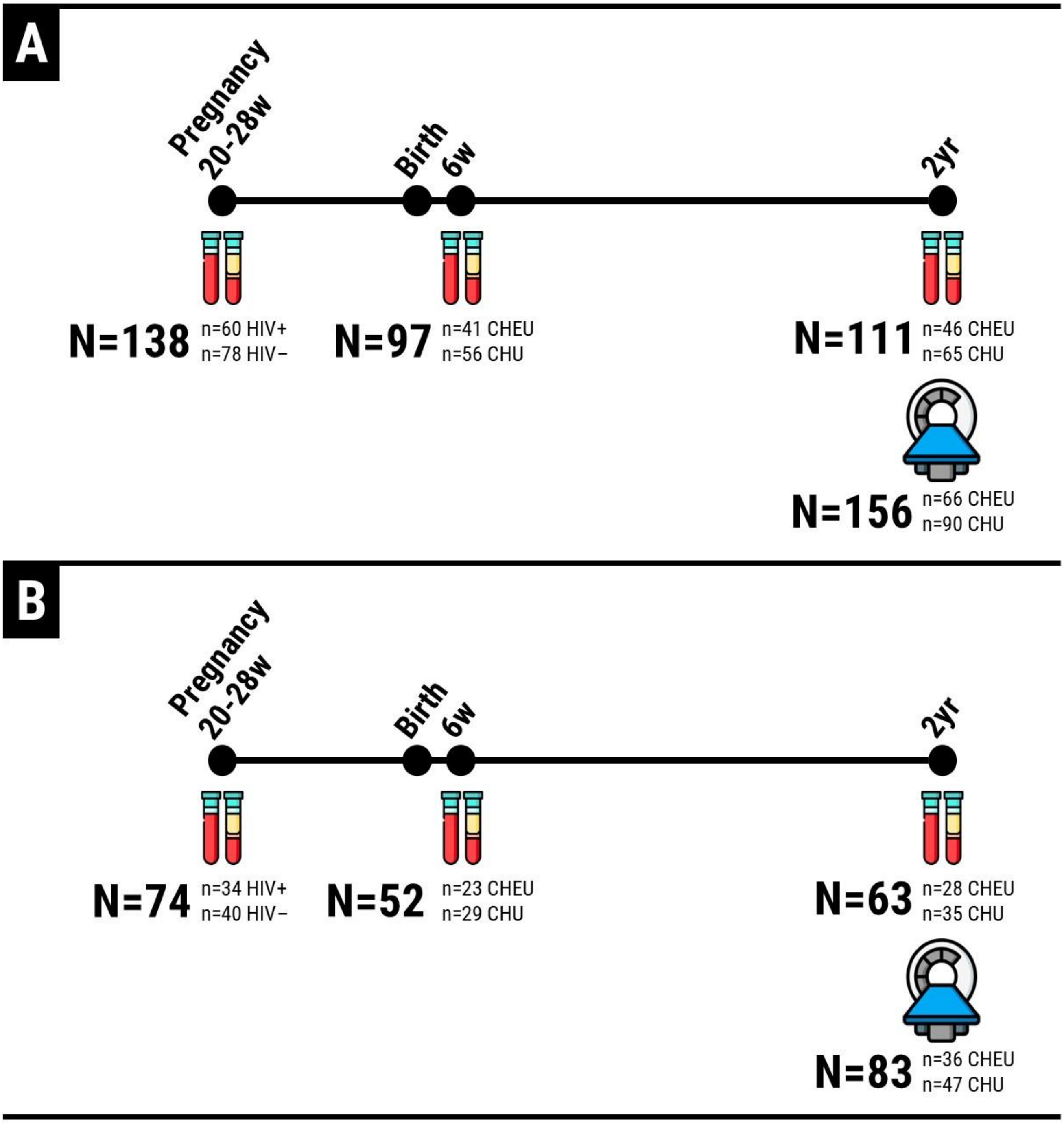

A. Number of participants with serum samples available at each timepoint, out of the entire cohort of children invited for magnetic resonance spectroscopy at 2-3 years of age.
B. Number of participants with serum samples available at each timepoint, out of the entire cohort of children with complete, high-quality magnetic resonance spectroscopy data at 2-3 years of age.

**CHEU**: Children who are HIV-Exposed and Uninfected; **CHU**: Children who are HIV-Unexposed.

9 **Supplementary Figure 2. Maternal, infant, and child serum marker concentrations**

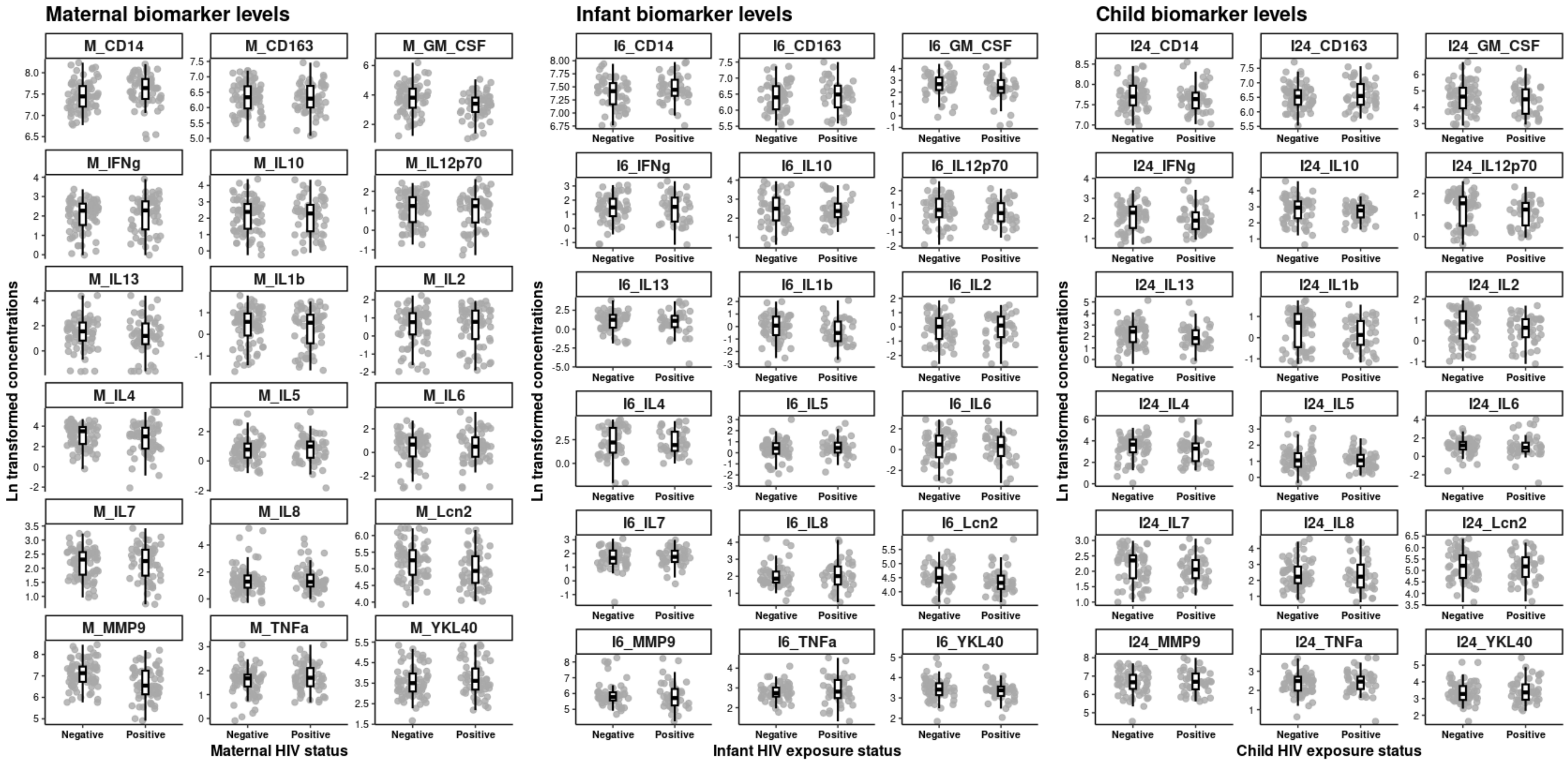

**Supplementary Figure 3. Trajectories of child serum marker concentrations from 6 weeks to 2 years of age**

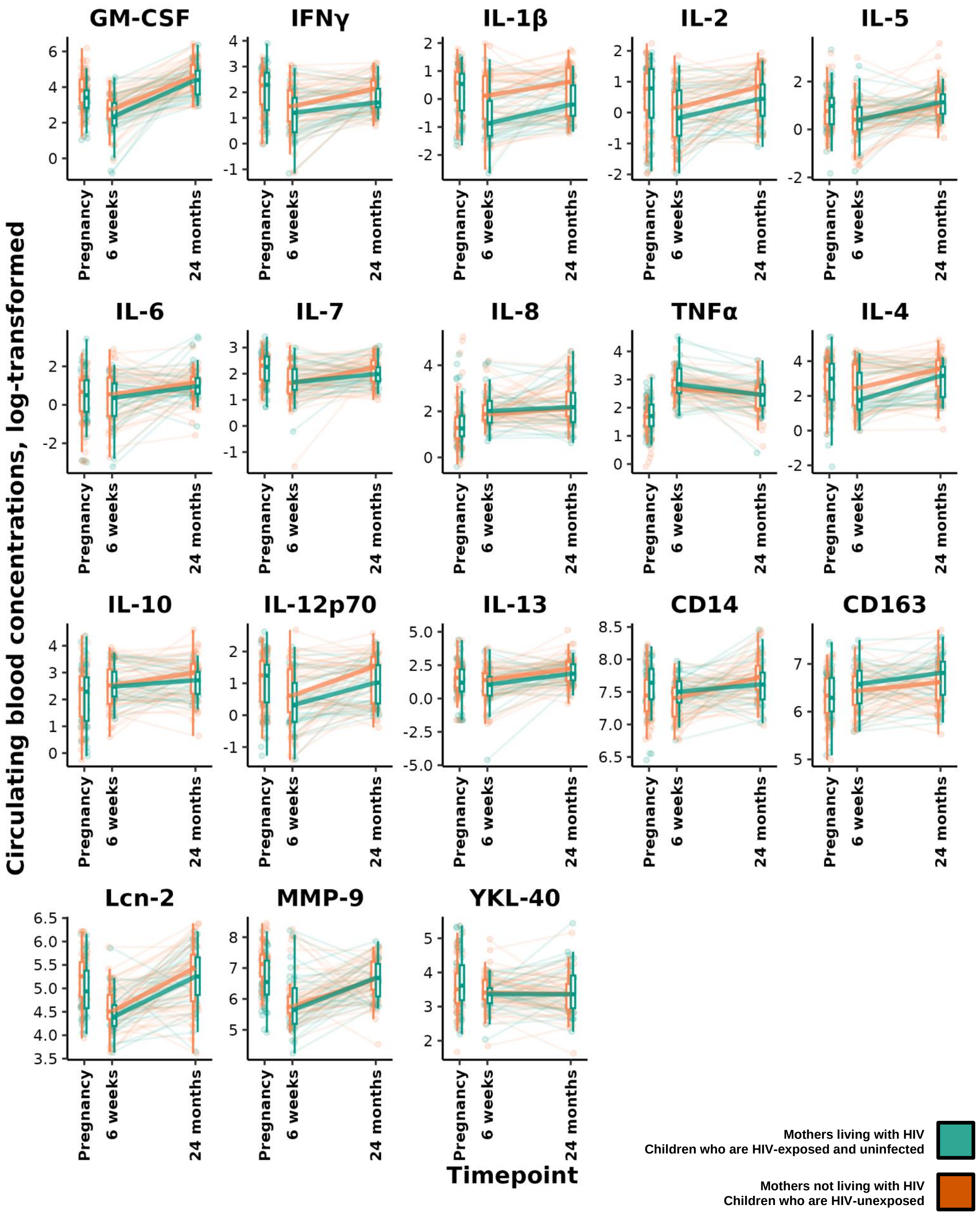

**Supplementary Figure 4. Forest plots: Associations between serum marker levels and child neurometabolite ratios in the HIV-exposed group**

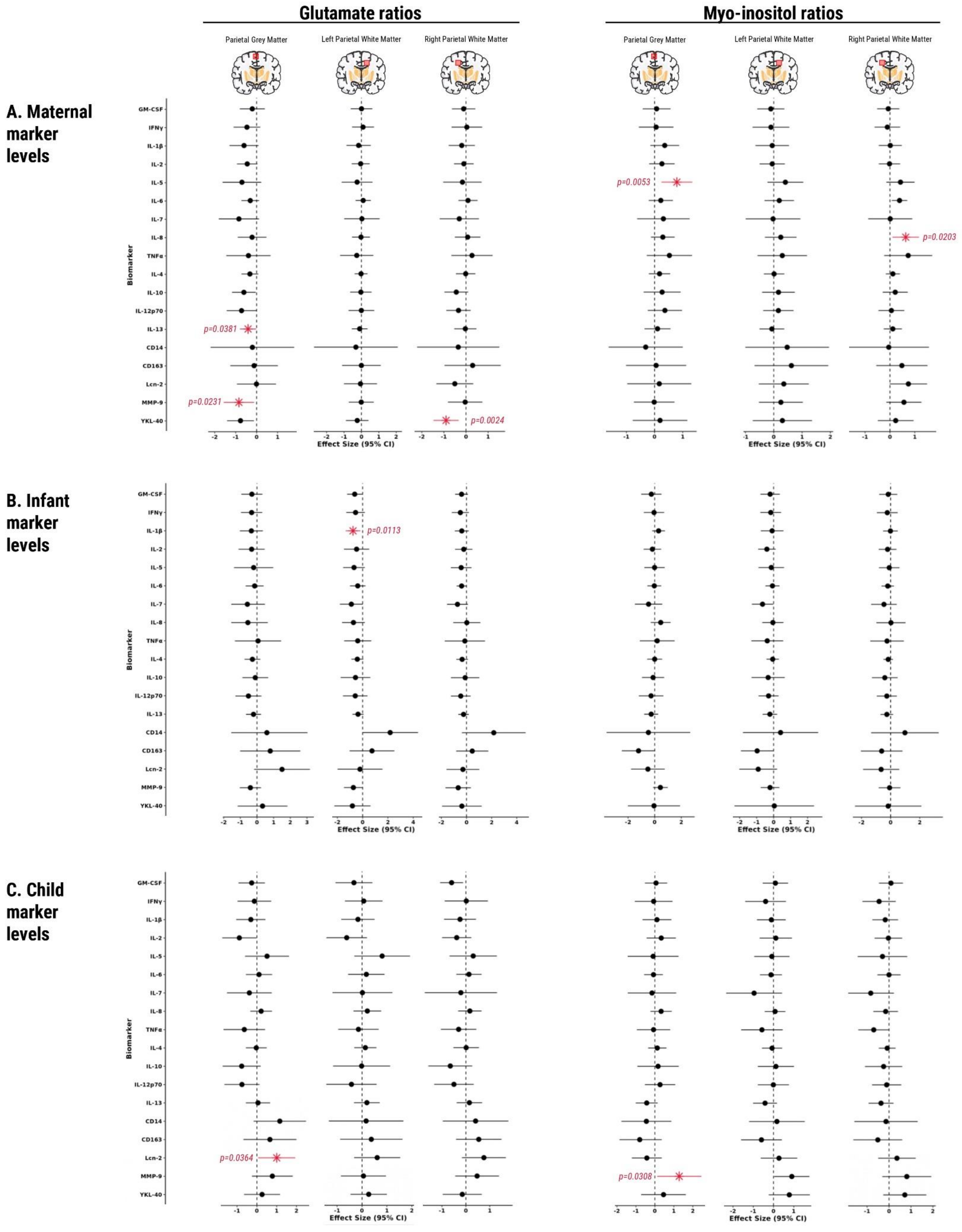
